# Unsupervised identification of the internal states that shape natural behavior

**DOI:** 10.1101/691196

**Authors:** Adam J. Calhoun, Jonathan W. Pillow, Mala Murthy

## Abstract

Internal states can shape stimulus responses and decision-making, but we lack methods to identify internal states and how they evolve over time. To address this gap, we have developed an unsupervised method to identify internal states from behavioral data, and have applied it to the study of a dynamic social interaction. During courtship, *Drosophila melanogaster* males pattern their songs using feedback cues from their partner. Our model uncovers three latent states underlying this behavior, and is able to predict the moment-to-moment variation in natural song patterning decisions. These distinct behavioral states correspond to different sensorimotor strategies, each of which is characterized by different mappings from feedback cues to song modes. Using the model, we show that a pair of neurons previously thought to be command neurons for song production are sufficient to drive switching between states. Our results reveal how animals compose behavior from previously unidentified internal states, a necessary step for quantitative descriptions of animal behavior that link environmental cues, internal needs, neuronal activity, and motor outputs.

## Introduction

Internal state can have a profound effect on behavioral decisions. For example, we are more likely to make correct choices when attending versus when distracted, and will consume food when hungry but suppress eating when sated. A number of studies in animals highlight that the nervous system encodes these context-dependent effects by remodeling sensorimotor activity at every level, from sensory processing, to decision-making, all the way to motor activity [1, 2, 6, 10, 14, 25, 43]. For instance, recordings from rodent cortical neurons reveal that neural activity can be more strongly correlated with the state of locomotion versus the statistics of sensory stimuli during sensory-driven tasks [50, 62]. Across model systems, locomotion can also change the gain of sensory neurons, causing them to be more or less responsive, as well as leading to production of distinct behavioral outputs when these neurons are activated [16, 31, 39, 47, 60, 63, 69]. And it is not simply action that modulates neural activity but also internal goals and needs. Circuits involved in driving courtship or aggression behaviors in rodents and flies show different patterns of activity, as motivation to court or fight changes [34, 42, 56, 74, 75]. Neurons can be modulated by multiple mechanisms in order to promote these goals: otherwise quiescent neurons can be activated. For example, during hunger states, chemosensory neurons that detect desirable stimuli are facilitated and enhance their response to these cues [45, 57, 66]. Downstream from sensory neurons, the needs of an animal can cause the same neurons to produce different behaviors - foraging instead of eating, for instance - when ensembles of neurons are excited or inhibited by neuromodulators that relay information about state [12, 22, 33, 35].

Despite evidence that internal states affect both behavior and sensory processing, we currently lack methods to identify the changing internal states of an animal over time. While some states can be controlled for or measured externally, such as nutritional status or walking speed, animals are also able to switch between internal states that are difficult to identify, measure, or control. One approach to solving this problem is to identify states in a manner that is agnostic to an animal’s sensory environment. These approaches attempt to identify whether the behavior an animal produces can be explained by some underlying discrete state, for example with a hidden Markov Model (HMM) [8, 27, 38, 71]. However, in many cases, the repertoire of behaviors produced by an animal may stay the same, while what changes is either the way in which sensory information patterns these behaviors or patterns the transitions between behaviors. Studies that dynamically predict behavior using past sensory experiences have provided important insight into sensorimotor processing but typically assume that an animal is in a single state [15, 19, 21, 64]. These techniques make use of regression methods such as generalized linear models (GLMs) that identify a ‘filter’ that describes how a given sensory cue is integrated over time to best predict future behavior. Here we take a novel approach to understanding behavior by using a combination of hidden state models (HMMs) and sensorimotor models (GLMs) to investigate the acoustic behaviors of the vinegar fly *D. melanogaster*.

Acoustic behaviors are particularly well-suited for testing models of state-dependent behavior. During courtship, males generate time-varying songs [5], whose structure can in part be predicted by dynamic changes in feedback cues, over timescales of 10s to 100s of milliseconds [17, 19, 20]; receptive females respond to attractive songs by reducing locomotor speed and eventually mating with suitable males [13, 18, 77]. Previous GLMs of male song structure did not predict song decisions across all courtship time [19]. We do so here, and find that by inferring hidden states, we can capture 84.6% of all remaining information about song patterning and 53% of all remaining information about transitions between song modes, both relative to a ‘Chance’ model that knows only about the distribution of song modes. This represents an increase of 70% for all song and 110% for song transitions over a GLM. The hidden states of the HMM rely on sensorimotor transformations, represented as GLMs, that govern not only the choice between song outputs during each state but also the probability of transitioning between states. Using GLM filters from the wrong state worsens predictive performance. We then use this model to identify neurons that induce state switching: we find that the neuron pIP10, previously identified as a part of the song motor pathway [70], additionally changes how the male uses feedback to modulate song choice. Our study highlights how unsupervised models that identify internal states can provide insight into nervous system function and the precise control of behavior.

## Results

### A combined generalized linear model (GLM) and hidden Markov model (HMM) effectively captures the variation in song production behavior

During courtship, *Drosophila* males sing to females in bouts composed of three distinct modes (Fig. 1a): ‘sine song’ and two types of ‘pulse song’ [17]. Previous studies used generalized linear models (GLMs) to predict song patterning choices (about whether and when to produce each song mode) from the feedback available to male flies during courtship [19, 20]; inputs to the models were male and female movements in addition to his changing distance and orientation to the female (Fig. 1b). This led to the discovery that males use fast-changing feedback from the female to shape their song patterns over time; moreover, these models identified the time course of cues derived from male and female trajectories that were predictive of song decisions. While these models could accurately predict up to of 60% of song patterning choices, when averaged across the data [19], they still left a large portion of variability unexplained.

**Figure 1.**
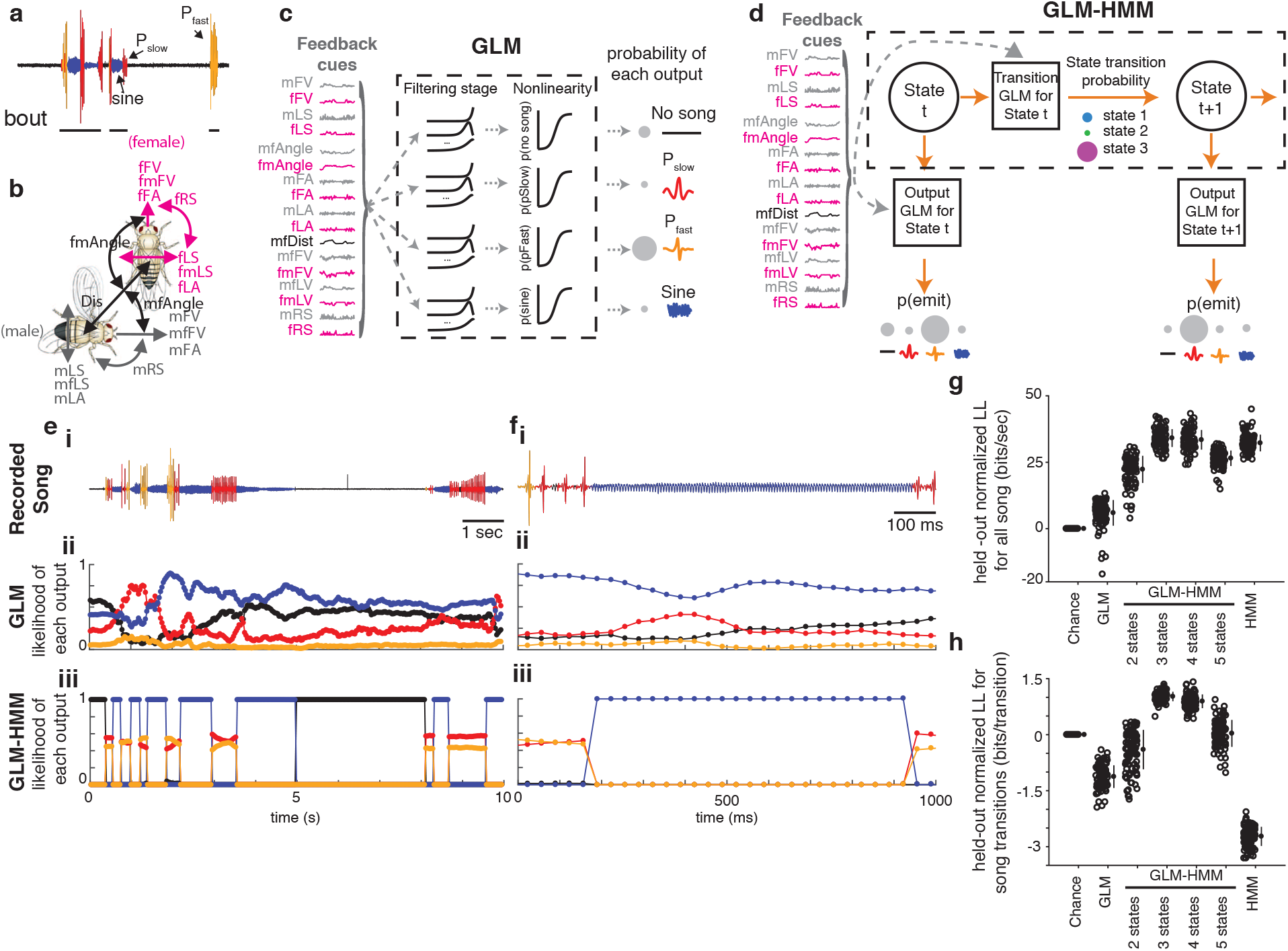
**a.** Fly song modes: No song (black), fast pulses or P_fast_ (orange), slow pulses or P_slow_ (red), and sine song (blue). Song is organized into trains of a particular type of song in a sequence (multiple pulses in a row constitute a pulse train) as well as bouts (multiple trains followed by no song, represented here by a black line). **b.** Fly feedback cues: male/female forward velocity (mFV/fFV), male/female lateral and rotational speeds (mLS/fLS and mRS/fRS), male/female forward and lateral accelerations (mFA/fFA and mLA/fLA), the component of male forward and lateral velocity in the direction of the female (mfFV and mfLS) and the component of the female forward and lateral velocity in the direction of the male (fmFV and fmLS), as well as the distance between the animals (mfDist) and the absolute angle from female/male heading to male/female center (mfAngle and fmAngle). **c.** Schematic illustrating the multinomial GLM which takes as input feedback cues and passes these through a linear filtering stage. There is a separate set of linear filters for each possible song mode. These filters are passed through a nonlinearity and the relative probability of observing each output (no song, P_fast_, P_slow_, sine song) gives the overall likelihood of song production. **d.** Schematic illustrating the GLM-HMM. At each time point t, the model is in a discrete hidden state. Each hidden state has a distinct set of multinomial GLMs that predict the type of song that is emitted as well as the probability of transitioning to a new state. **e.** i, Ten seconds of natural courtship song consisting of no song (black), P_fast_ (orange), P_slow_ (red), and sine (blue). ii, The conditional probability of each output type for this stretch of song under the standard GLM. iii, The conditional probability of the same song data under the 3-state GLM-HMM; predictions are made one step forward at a time using past feedback cues and song mode history (see Methods). **f.** (i), One second of natural song. (ii) and (iii), conditional probability of each song mode under GLM (ii) and GLM-HMM (iii), as in (e). **g.** Normalized log-likelihood on test data (in bits/sec; see Methods). The GLM outperforms the ‘Chance’ model (p < 1e-20), but the 3-state GLM-HMM produces the best performance (each open circle represents predictions from one courtship pair (only 100 of the 276 pairs shown for visual clarity); filled circles represent mean +/− SD). The 3-state model outperformed a 2-state GLM-HMM (p < 1e-40) and the 5-state GLM-HMM (p < 1e-40), but is not significantly different from a 4-state model (p = 0.18). By this metric, the 3-state GLM-HMM slightly outperforms a HMM (1 bit/sec improvement, p < .001). All p-values from two-tailed t test. **h.** Normalized test log-likelihood during transitions between song modes (e.g., transition from sine to P_slow_). The 3-state GLM-HMM outperforms the GLM (p < 1e-100), the 2-state GLM-HMM (p < 1e-60), and substantially outperforms the HMM (p < 1e-160).

Leveraging a previously collected data set of 2,765 minutes of courtship interactions from 276 wild type pairs [19], we trained a multinomial GLM [28] (Fig. 1c) to predict song behavior over all courtship time (we predict four song modes: sine, fast pulses (P_fast_), slow pulses (P_slow_), and no song) (Fig. 1a) from the time histories of 17 potential feedback cues defined by male and female movements and interactions (Fig. 1b) - what we will refer to as ‘feedback cues’. The overall prediction of this model (Fig. 1e-h, “GLM”) was similar to prior work [19] which used a smaller set of both feedback cues and song modes, and a different modeling framework (Supplemental Fig. 1a-b, (compare “Coen 2014” with “This study” and “GLM”)). We compared this model to one in which we examined only the mean probability of observing each song mode across all of courtship (“Chance”).

We next created a model that incorporates hidden states (also known as latent states) when predicting song from feedback cues, a model we term the GLM-HMM (generalized linear model - hidden Markov model; Fig. 1d). A standard HMM has fixed probabilities of transitioning from one state to another, as well as fixed probabilities of emitting different actions in each state. The GLM-HMM allows each state to have an associated multinomial GLM to describe the mapping from feedback cues to the probability of emitting a particular action (one of three types of song or no song). Each state also has a multinomial GLM that produces a mapping from feedback cues to the transition probabilities from the current state to the next state (Supplemental Fig. 1c-d). This allows the probabilities to change from moment to moment in a manner that depends on the feedback the male receives, and to determine which feedback cues affect the probabilities at each moment. This model is inspired by previous work modeling neural activity [26], but uses multinomial categorical outputs to account for the discrete nature of male singing behavior. One major difference between the GLM-HMM and other models that predict behavior [9, 61, 71] is that our model allows each state to predict behavioral outputs with a different set of regression weights.

We used the GLM-HMM to predict song behavior (Supplemental Fig. 2a-b) and compared its predictive performance on held out data to a ‘Chance’ model, which only captures the marginal distribution over song modes (e.g., males produce ‘no song’ 68% of the time). We quantified model performance using the difference between model log-likelihood and log-likelihood of the ‘Chance’ model, normalized to have units of bits/sec (see Methods). While the GLM has no internal states to estimate, for all models with an HMM, we used the feedback cue and song mode history to estimate the probability of being in each state. We then predicted the song mode in the next time bin using this probability distribution over states (see Methods). For predicting all song, or every bin across held out data, a 3-state GLM-HMM outperforms the GLM (Fig. 1e-f (compare middle and bottom rows)). We found an improvement of 32 bits/sec relative to the ‘Chance’ model for the 3-state GLM-HMM (versus 6 bits/sec for the GLM). We binned song data into 30 bins/sec, so a performance of 60 bits/sec would indicate that our model has the information to predict every bin with 100% accuracy. Knowledge of the mean song probability (the ‘Chance’ model) reduces the uncertainty to 40.3 bits/sec. The 3-state GLM-HMM, therefore, captures 84.6% of the remaining information, as opposed to only 14.6% with a GLM (Fig. 1g). Fitting models with additional states did not significantly improve performance, and tended to decrease predictive power, likely due to over-fitting. The 3-state GLM-HMM also offered a significant improvement in prediction over ‘Chance’, even when only the history of feedback cues (not song mode history) was used for predictions (Supplemental Fig. 2e-f; see Methods). However, an HMM is nearly as good at predicting all song, because the HMM largely predicts in the next time bin what occurred in the previous time bin (see Methods), and song consists of runs of each song mode (Fig. 1g; although this depends heavily on the autocorrelation structure of the binned data (Supplemental Fig. 2c)). A stronger test then is to examine song transitions (for example, times at which the male transitions from sine to P_fast_ or P_slow_ to no song) — in other words, predicting when the male changes what he sings. The 3-state GLM-HMM offers 1.02 bits/transition over a ‘Chance’ model (capturing 53% of the remaining information about song transitions), while the HMM is significantly worse than the ‘Chance’ model at −2.7 bits/transition (Fig. 1h and Supplemental Fig. 2d). In fact, the 3-state GLM-HMM is substantially better than all other models at predicting song transitions (an increase of 110% over the GLM, for example) — such events are rare, and therefore not well captured when examining the performance across all song. Moreover, the three state GLM-HMM outperformed previous models [19], even considering that those models were fitted to subsets of courtship data (for example, only times when the male was oriented towards the female), effectively binning the data into particular ‘contexts’ (Supplemental Fig. 1b). Thus, the GLM-HMM can account for much of the moment-to-moment variation in song patterning by allowing for three distinct sensorimotor strategies or states. We next investigated what these states correspond to and how they impact behavior.

### Three distinct sensorimotor strategies during song production

We determined how the 17 feedback cues and 4 song modes differed across the three states of the GLM-HMM. We examined mean feedback cues (Fig. 1b) during each state. We found that in the first state, the male is closer to the female on average and moving slowly in the direction the female we therefore term this state the ‘close’ state (Fig. 2a, Supplemental Fig. 3a,d). In the second state, the male is, on average, moving toward the female at higher speed, while still at a relatively close distance, and so we call this the ‘chasing’ state (Fig. 2b, Supplemental Fig. 3b,e). In the third state, the male is, on average, further from the female, moving slowly, and oriented away from the female, and so we call this the ‘whatever’ state (Fig. 2c, Supplemental Fig. 3c,f). However, there is also substantial overlap in the distribution of feedback cues that describe the average of each state (Fig. 2d-g), indicating that the distinction between each state is more than just these descriptors. Another major difference between the states is the song output that dominates in each state - the ‘close’ state mostly generates sine song, while the ‘chasing’ state mostly generates pulse song, and the ‘whatever’ state mostly no song (Fig. 2a-c). However, we note that there is not a simple one-to-one mapping between states and song outputs [36]. All 4 outputs (no song, P_fast_, P_slow_, and sine) are emitted in all three states, and the probability of observing each output depends on the feedback cues the animal receives at that moment.We compared this model to a GLM-HMM with four states, which performs nearly as well as the 3-state GLM-HMM (Fig. 1g-h). We found that three of the four states correspond closely to the 3-state model (Supplemental Fig. 3g-h). When animals are in the fourth state of the 4-state model, they are mostly in the ‘whatever’ state of the 3-state model - but enter this state less than 1% of the time (Supplemental Fig. 3i). We conclude that the 3-state model is the most parsimonious description of *Drosophila* song patterning behavior.

**Figure 2.**
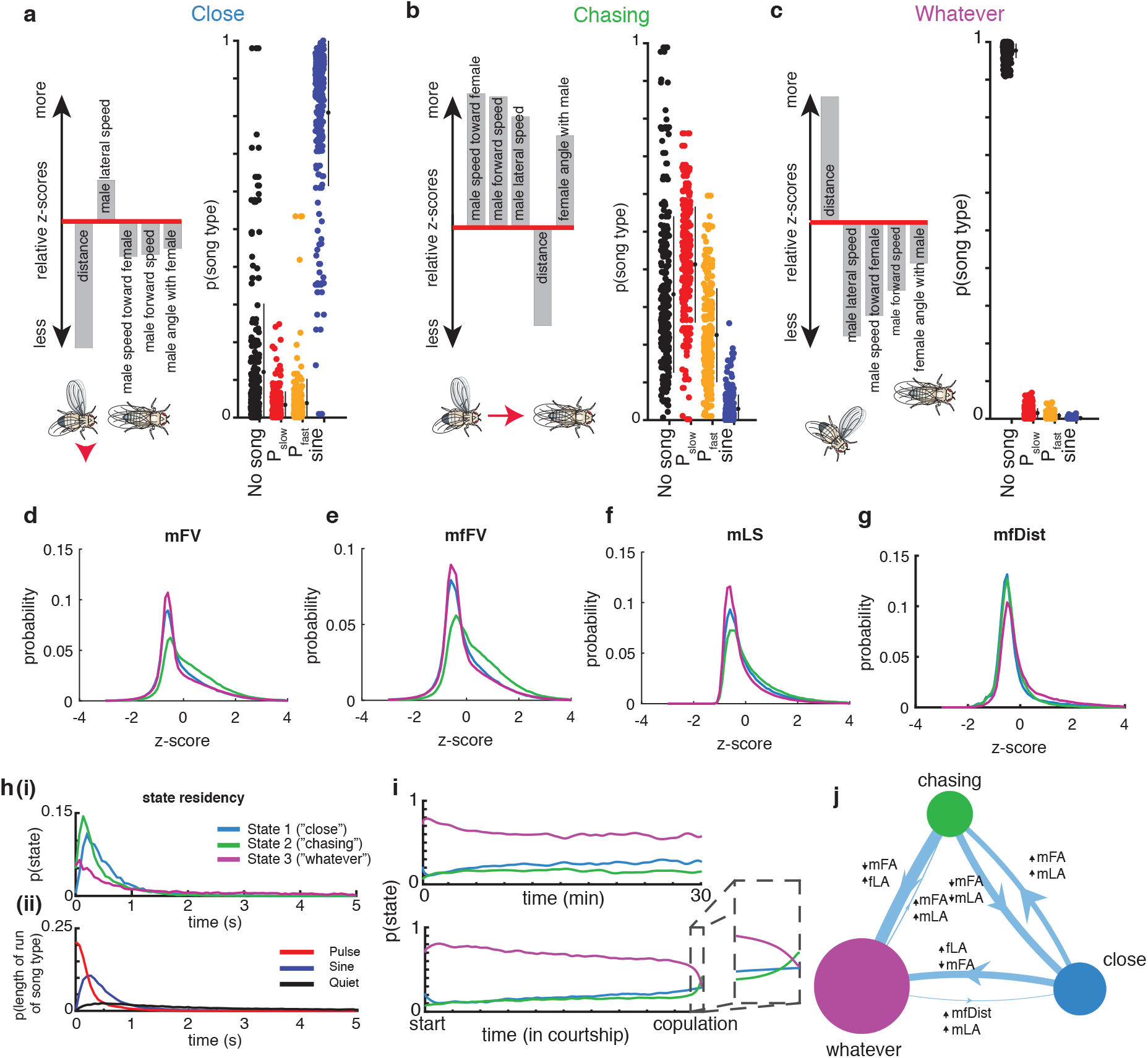
**a-c**. Left panels, the five feedback cues that are most different from the mean when the animal is in the **(a)** ‘close’, **(b)** ‘chasing’, or **(c)** ‘whatever’ state (see Methods for details on z-scoring). Illustration of flies is a representation of each state according to these feedback cues. Right panels, The probability of observing each type of song when the animal is in that state. Filled circles represent individual animals (n=276 animals, small black circles with lines are mean +/− SD). **d-g**, Distributions of values (z-scored, see Methods) for four of the feedback cues (see Fig. 1b) and for each state. Although a state may have features that are larger or smaller than average, the distributions are highly overlapping. **h. (i)** The dwell times of the ‘close’, ‘chasing’, and ‘whatever’ states across all of the data (including both training and validation sets). **(ii)** The dwell times of sine trains, pulse trains, and stretches of no song (see Fig. 1a for definition of song modes; P_fast_ and P_slow_ are grouped together here) across all of the data are dissimilar from the dwell times of the states with which they are most associated. **i. (i)**. The mean probability across flies of being in each state fluctuates only slightly over time when aligned to absolute time (top) or the time of copulation (bottom). Just prior to copulation, there is a slight increased probability of being in the ‘chasing’ state (ii, zoom). **j**. Areas of circles represent the mean probability of being in each state and width of each line represents the fixed probability of transitioning from one state to another. The filters that best predict transitioning between states (and modify the transition probabilities) label each line, with the ‘up arrow’ representing feedback cues that increase the probability and ‘down arrow’ feedback cues that decrease the probability.

Hidden Markov models are memoryless; the past has no influence on which state the model will next enter, and thus these models exhibit dwell times that follow exponential distributions. However, natural behavior exhibits very different distributions [7, 61, 71]. In our model, which is no longer stationary (memoryless), we find that the dwell times in each state follow a non-exponential distribution (Fig. 2h(i), Supplemental Fig. 3j); moreover, the majority of dwell times are on the order of hundreds of milliseconds to a few seconds, indicating that males switch between states even within song bouts (Fig. 2h(ii)). It is possible that states correspond to differences in behavior that change progressively throughout courtship - to test this, we examined the mean probability across animals of males being in each of the three states across absolute time (Fig. 2i, top) or aligned to successful copulation (Fig. 2i, bottom). We find that the probability of being in each of the three states is steady throughout courtship, without a clear trend towards one or more states over time. The only exception is that just prior to copulation males are more likely to sing and, accordingly, the probability of the ‘chasing’ state increases, while the probability of the ‘whatever’ state decreases (Fig. 2i inset) — though note that this does not mean that the animals were not physically close to the female (Fig. 2g). We next examined which feedback cues (Fig. 1b) predict the transitions between states (Fig. 2j); we refer to these GLM filters as state-transition filters. Interestingly, the feedback cues that are most predictive of transitions between states (Fig. 2j, Supplemental Fig. 4a-c) are different from the feedback cues with the largest magnitude mean value in each state (Fig. 2a-c), suggesting that the dynamics of what drives an animal out of a state is different from the dynamics that are ongoing during production of a state.

### Feedback cues possess different relationships to song behavior in each state

The fact that each song mode (no song, P_fast_, P_slow_, and sine) is produced in each state of the 3-state GLM-HMM (Fig. 2a-c) suggests that the difference between each state is not the type of song that is produced but the GLM filters that predict the output of each state (which we will refer to as the ‘output filters’). To test this hypothesis we generated discretized song based on either the full GLM-HMM model, or using output filters from only one of the three states (Fig. 3a). This confirmed two features of the model. First, that each set of output filters can predict all possible song outputs (no song, P_fast_, P_slow_, and sine) depending on the input. Second, the song prediction from any one state is insufficient for capturing the overall moment-to-moment changes in song patterning. To quantify this, we performed a similar analysis over 926 minutes of courtship data: we determined how well the model would predict each song behavior (no song, P_fast_, P_slow_, and sine), we asked what the conditional probability was of observing that song either using only the output filters of that state or only using the output filters of one of the other states (Fig. 3b). We found that the predictions were highly state-specific and performance degrades dramatically when using filters from the wrong state. For example, even though the ‘close’ state is mostly associated with the production of sine song (Fig. 2a), the production of both types of pulse song and no song during the ‘close’ state are best predicted using the output filters from the ‘close’ state. The same holds for the ‘chasing’ and ‘whatever’ states — the dynamic patterning by feedback cues shows the distinction between states is not merely based on the different types of song. Taken together, the highly divergent predictions by state (Fig. 3a) and the lack of explanatory power from other states’ output filters (Fig. 3b) suggest that ‘close’, ‘chasing’, and ‘whatever’ are in fact distinct states.

**Figure 3.**
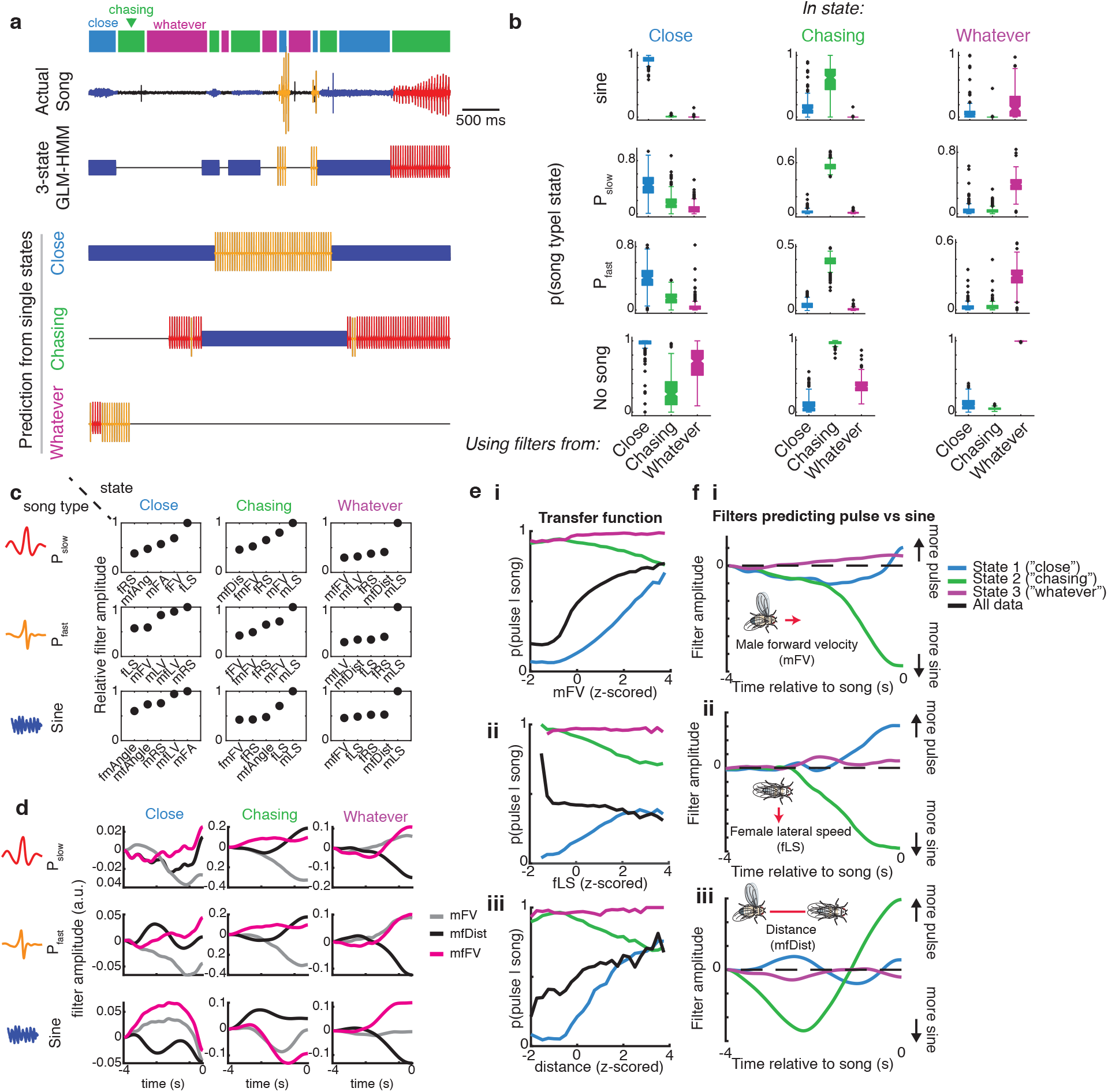
**a.** A stretch of 5 seconds of song production from the natural courtship data set, with the prediction of states indicated above in colored squares (blue = ‘close’, green = ‘chasing’, and purple = ‘whatever’). The prediction of the full GLM-HMM model (third row) is very different from the prediction if we assume the animal is always in the ‘close’ state, ‘chasing’ state, or ‘whatever’ state. The output using the song prediction filters from only that state is illustrated in the bottom three rows. **b.** The conditional probability (across all data, n=276 animals, error bars represent SEM) of observing a song mode in each state (predicted by the full 3-state GLM-HMM), but using output filters from only one of the states. Conditional probability of the appropriate state is larger than the conditional probability of the out-of-state prediction (p < 1e-4 for all comparisons, one-sample, one-tailed t test corrected with Bonferroni correction). Song mode predictions are highest when using output filters from the correct state. Center lines of box plots represent median, the bottom and top edges represent the 25th and 75th percentiles (respectively). Whiskers extend to +/− 2.7 times the standard deviation. **c.** The five most predictive output filters for each state and for prediction of each of three of the types of song. Filters for types of song are relative to ‘no song’ filters which are set to a constant term (see Methods). **d.** Example output filters for each state reveal that even for the same feedback cues, the GLM-HMM shows distinct patterns of integration. Plotted here, male forward velocity (mFV), distance (mfDist), and the male’s velocity in the direction of the female (mfFV) - filters can change sign and shape between states. **e.** Transfer functions (the conditional probability of observing song choice (y axis) as a function of the magnitude of each feedback cue (x axis)) for producing pulse (both P_slow_ and P_fast_) versus sine have distinct patterns based on state. For **(i)** male forward velocity, **(ii)** female lateral speed, and **(iii)** distance, the average relationship or transfer function between song choice and the movement cue (black) differs with transfer functions separated by state (blue, green, and purple). **f.** Output filters that predict pulse versus sine song, for each of three feedback cues: **(i)** male forward velocity, **(ii)** female lateral speed, and **(iii)** distance.

Which feedback cues (Fig. 1b) are most predictive of song decisions in each state? To answer this question, we examined all predictive feedback cues of each song type in each state, in rank order (Supplemental Fig. 5; Fig. 3c plots the top 5 most predictive feedback cues per state and per song type). This revealed that the most predictive feedback cues are strongly re-weighted by state - for instance, male lateral velocity is the largest predictor of both P_fast_ and P_slow_ in the ‘chasing’ and ‘whatever’ states but is not one of the top 5 predictors in the ‘close’ state. As another example, female lateral speed is most predictive of P_slow_ in the ‘close’ state, but is not in the top 5 predictors for the ‘chasing’ state. By comparing the output filters of the same feedback cues for different states, we observed that both the temporal dynamics (Supplemental Fig. 6a-c) and the sign of the output filters can change by state (Supplemental Fig. 6d-f). Taken together, these observations suggest that state switching occurs due to both re-weighting of which feedback cues are important as well as re-shaping of the output filters themselves.

We next sought to identify whether the state-specific sensorimotor transformations uncovered relationships between feedback cues (Fig. 1b) and song behaviors that were previously hidden. Previous work on song patterning identified male forward velocity (mFV), female lateral speed (fLS), and male-female distance (mfDist) as important predictors of song structure [19], with increases in mFV and mLS predicting pulse song, while decreases predicted sine song; fLS was most predictive of the end of song bouts. By contrast, our model reveals that, while on average (and across all states) the amount of pulse song increases with male velocity (Fig. 3e(i), black), this was only true for the ‘close’ state, and the relationship (or transfer function) was actually inverted in the ‘chasing’ state (Fig 3e(i), green and blue). Increased female lateral speed was previously shown to increase the probability of switching song type from pulse to sine song [19]; however, when we examine fLS by state, we found this feature is positively correlated with the production of pulse song in the ‘close’ state (Fig. 3e(ii), blue) but negatively correlated with the production of pulse song in the ‘chasing’ state (Fig. 3e(ii), green). Finally, the distance between males and females was previously shown to predict the choice to sing pulse (at greater distances) over sine (at shorter distances) - the relatively quieter sine song is produced when males and females are in close proximity [17]. Again, we found this to be true only when the animals are in the ‘close’ state (Fig. 3e(iii)) but the relationship between distance and song type (pulse versus sine) is inverted in the ‘chasing’ state. Interestingly, when we examine the feedback cue filters (Fig. 3f) we find that while mFV and fLS are cumulatively summed to predict song type, the distance filter is a long-timescale differentiator across different timescales in each state, as opposed to the short-timescale integrator found previously [19]. Our GLM-HMM therefore reveals unique relationships between input and output that were not uncovered when data was aggregated across states.

### Activation of pIP10 neurons biases males towards the ‘close’ state

Having uncovered three distinct sensorimotor patterning strategies via the GLM-HMM, we next used the model to identify neurons that modulate state-switching. To do this, we optogenetically activated candidate neurons that might be involved in driving changes in state specifically during acoustic communication; we reasoned that such a neuron might have already been identified as part of the song motor pathway [34, 41, 70]. The goal was to perturb the circuitry underlying state-switching, and thereby change the mapping between feedback cues and song modes. We focused on three classes of neurons that when activated produce song in solitary males: P1a, a cluster of neurons in the central brain, pIP10, a pair of descending neurons, and vPR6, a cluster of ventral nerve cord (VNC) pre-motor neurons (Fig. 4a). Across a range of optogenetic stimulus intensities, P1a and pIP10 activation in solitary males induces the production of all three (P_fast_, P_slow_, and sine) types of song, whereas vPR6 activation induces only pulse song (P_fast_ and P_slow_) [17]. We hypothesized that activation of these neurons could produce changes in song either through directly activating motor pathways or through changing the transformation between sensory information and motor output. Previous work demonstrated that visual information related to estimating the distance between animals is likely relayed to the song pathway between pIP10 neurons and VNC song pre-motor neurons [20]. pIP10 neurons could therefore influence how sensory information modulates the song pre-motor network, and consequently affect the mapping between feedback cues and song modes.

**Figure 4.**
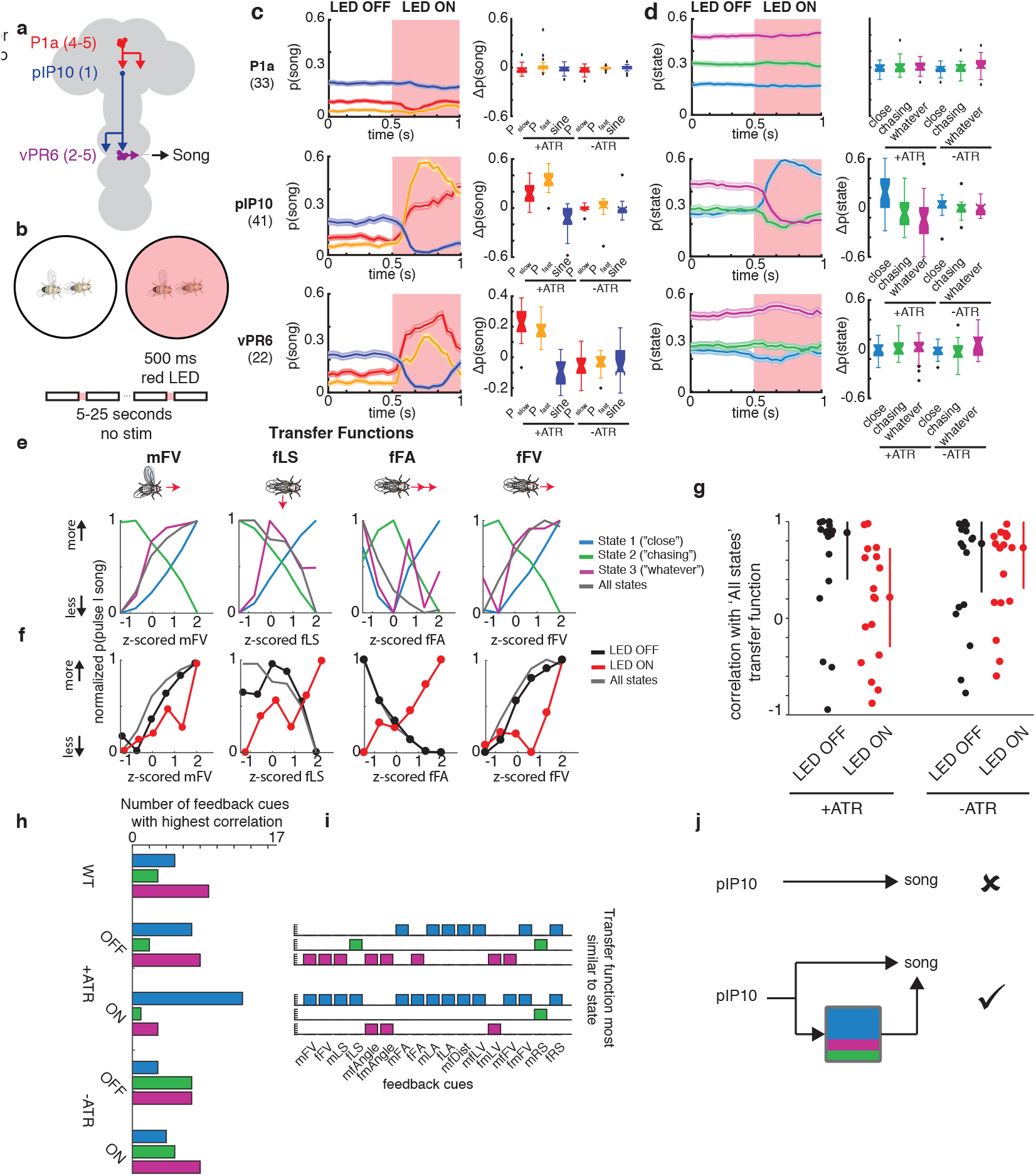
**a.** Schematic of 3 classes of neurons in the *Drosophila* song production pathway. **b.** Protocol for optogenetically activating song pathway neurons using csChrimson targeted to each of the neuron types in (a). **c.** The observed probability of each song mode aligned to the onset of the optogenetic stimulus (left). Difference between mean during LED on from mean during LED off prior to stimulation (right). Number of flies tested are indicated in parentheses; error bars are SEM. Control males are of the same genotype but have not been fed all-trans retinal (ATR), the required co-factor for csChrimson. **d.** The posterior probability of each state given the feedback cues and observed song (under the 3-state GLM-HMM trained on wild type data), aligned to the onset of optogenetic stimulation (left). Activation of pIP10 neurons biases males to the ‘close’ state and away from the ‘chasing’ and ‘whatever’ states. (right) Difference between mean during LED on from mean during LED off prior to stimulation (right); error bars are SEM. **e.** Comparison of transfer functions (the conditional probability of observing song choice (y axis) as a function of the magnitude of each feedback cue (x axis)); (see Fig. 3e). Shown here are transfer functions for four feedback cues (male forward velocity (mFV), female lateral speed (fLS), female forward acceleration (fFA), and female forward velocity (fFV). Average across all states (dark gray) represents the transfer function from all data without regard to the state assigned by the model.Transfer functions are calculated from all data. **f.** Transfer functions for the same four feedback cues shown in (e), but in animals expressing csChrimson in pIP10 while the LED is off (black) or on (red); transfer functions for data from wild type animals across all states (dark grey) reproduced from (e). **g.** For all seventeen feedback cues, median correlation between transfer functions between ‘all states’ and the four conditions (pIP10 with ATR+ (LED off or on) or ATR− (LED off or on). Error bars represent median absolute deviation. **h.** The number of feedback cues with the highest correlation between the wild type transfer functions (separated by state) and the transfer functions for each of the conditions (pIP10 + ATR (LED off or on) and pIP10 ATR− (LED off or on). Blue represents transfer functions most similar to the ‘close’ state, green to the ‘chasing’ state, and purple to the ‘whatever’ state. **i.** Unpacking the data in (h) for the +ATR condition. **j.** (top) Previous view of pIP10 function. (bottom) Here, we show that pIP10 activation both drives song production and state-switching – this revised view of pIP10 function would not have been possible without the computational model.

We expressed the light-sensitive opsin csChrimson [40] using driver lines targeting P1a, vPR6 and pIP10, and chose a light intensity level, duration, and interstimulus-interval that reliably produces song in solitary males for these genotypes [17]; here, we activated neurons in the presence of a female, with varying pauses between stimuli to induce a change in state without completely overriding the role of feedback cues (Fig. 4b). We recorded song via an array of microphones tiling the floor of the chamber and wrote new software, called DeepFlyTrack, for tracking the centroids of flies on such difficult image backgrounds (see Methods). For this stimulation protocol, P1a drove a general increase in song during courtship (Supplemental Fig. 7a-c) while pIP10 and vPR6 activation reliably drove song production during the optogenetic stimulus (Fig. 4c, Supplemental Fig. 7c). The activation of P1a during courtship was different from previous findings in solitary males which showed stimulus-locked changes in song production [17] (Supplemental Fig. 7b), while type and quantity of song production from pIP10 and vPR6 activation was more similar [17]. To determine whether optogenetic activation affected state switching, we fit our GLM-HMM to recordings from males of all three genotypes (including both experimental animals and controls not fed the csChrimson channel co-factor, all trans retinal (‘ATR-’) - see Methods).

To account for the possibility that activating these neurons directly drives song production, we supplemented the GLM-HMM model (Fig. 1) with a filter encoding the presence or absence of the optogenetic LED stimulus. This filter (termed the ‘opto filter’) was fit separately for each genotype (see Table 1) and accounts for the change in probability of producing song that is unrelated to sensory information. This allowed us to directly compare the states found in wild type flies to optogenetically activated flies. The opto filters for each output type were similar across all states in ATR-fed flies (Supplemental Fig. 7e-f), indicating that any differences we found between states could be attributable to other aspects of the model (not the presence or absence of the LED light/optogenetic stimulus). Flies not fed ATR had opto filters that showed no influence of the LED stimulus on song, as expected (Supplemental Fig. 7f). We found that there was a large increase in the probability of entering the ‘close’ state when pIP10 neurons were activated, but little effect on state when P1a or vPR6 neurons were activated (Fig. 4d, Supplemental Fig. 7d,g). We found a consistent effect when we tested another line that labeled pIP10 [23] (data not shown). The ‘close’ state is typically associated with an abundance of sine song, though it also produces all other song modes during natural behavior (Fig. 2a); nonetheless, in this case, pIP10 activation was associated with increased pulse song (Fig. 4c; the opto filters predict P_fast_ and P_slow_ song for this manipulation (Supplemental Figure 7e)). Even though the male mostly sings pulse song during optogenetic activation of pIP10, the dynamics of the feedback cues that predict that pulse song are better matched to the output filters of the ‘close’ state of the GLM-HMM. Activation of pIP10 neurons always increased the probability that the animal would transition into the ‘close’ state, independent of which state the animal was in prior (however if the male was already in the ‘close’ state, there was no significant change in state (Supplemental Fig. 8a)), whether the animal was close to or far away from the female (Supplemental Fig. 8b), or singing or not singing (Supplemental Fig. 8c).

We next explored the possibility that the effect was somehow due to nuances of model fitting. Because vPR6 activation results in changes in song that are similar in aggregate to pIP10 activation (Fig. 4c) without a change in state (Fig. 4d), we conclude that changing state is not synonymous with changing song production. In addition, we found that the change in state could not be simply explained by pIP10 activation having a direct effect on both male and female movements and song (Supplemental Fig. 9a-d). We removed male and female feedback cues from the GLM-HMM by zeroing out their values (see Methods) and found that a model without feedback cues was poor at predicting song choice, suggesting that the prediction of ‘close’ state relies on the moment-to-moment variation in these features (Supplemental Fig. 9a). In addition, we examined individual feedback cues to see whether the change in state was due to a change in only a few feedback cues. We found that the vast majority of the feedback cues are more like the ‘close’ state (Supplemental Fig. 9b-c). We finally asked whether activation of pIP10 neurons puts the animal in a different behavioral context with respect to the female (e.g., driving him closer to the female). By looking at the six feedback cues (before and during optogenetic stimulation) that were the strongest predictors of being in the ‘close’ state, we found the dynamics of most feedback cues were indistinguishable from ATR- controls and could not explain the observed difference in state (Supplemental Fig. 9d). Instead, our data point to the fact that pIP10 neurons affect the way in which feedback cues (Fig. 1b) modulate song choice.

We next asked whether we could observe, following pIP10 activation, a change in song strategy independent of the GLM-HMM model. This was to ensure that our results were not simply due to how we fit the model, but rather could be observed in the data alone. As in Fig. 3e, we examined the male’s choice between producing pulse versus sine song. Because pIP10 directly drives an overall increase in pulse song (Fig. 4c), we normalized the data to the highest and lowest pulse rate, to more easily visualize the transfer function (the relationship between feedback cues and song choice) used by the male (Fig 4e). The relationship between feedback cues and the probability of producing pulse song is reversed between the ‘close’ and the ‘chasing’ state. We examined the correlation between these transfer functions from wild type males and males with pIP10 activation during LED ON or LED OFF (see Methods). We found that these transfer functions are similar to wild type (combining across all 3 states) for pIP10 activated flies during LED OFF, but are highly dissimilar when the LED is ON, suggesting that song patterning changes during these times (Fig 4f-g, Supplemental Fig. 10a). We then asked whether these transfer functions are shifted in a particular direction, such as toward the functions from the ‘close’ state in the wild type data (Supplemental Fig. 10b, blue lines). During periods when the LED is OFF, the transfer function resembles a mix of states (Fig. 4h). However, transfer functions during LED ON shift towards ‘close’ state transfer functions (Fig 4h). This was true across 13/17 feedback cues (Fig. 4i and Supplemental Fig. 10d-f). This analysis, independent of the GLM-HMM, confirms that pIP10 activation biases the nervous system to the ‘close state’ set of sensorimotor transformations that shape song output. pIP10 neurons therefore play a dual role in the acoustic communication circuit during courtship (Fig. 4j) – they both directly drive pulse song production (Fig. 4c) and also bias males towards the ‘close’ state (Fig. 4i). These results highlight the value of the GLM-HMM for identifying the neurons that influence dynamically-changing internal states, and are critical for shaping behavior.

## Discussion

Here we develop a model (the GLM-HMM) that allows experimenters to identify, in an unsupervised manner, dynamically changing internal states that influence decision-making, and ultimately, behavior. Using this model, we found that during courtship, *Drosophila* males utilize three distinct sensorimotor strategies (the 3 states of the model). Each strategy corresponds to a different relationship between inputs (17 feedback cues that affect male singing behavior) and outputs (3 types of song and no song). While previous work revealed that fly feedback cues predict song patterning decisions [17, 19], the discovery of distinct state-dependent sensorimotor strategies was only possible with the GLM-HMM. This represents an increase of 70% for all song and 110% for song transitions over a GLM. While we have here accounted for much of the variability in song patterning, we speculate that the remaining variability is due to either some noise in our segmentation of song [3] or the fact that we do not measure some male behaviors that are known to be part of the courtship interaction, including tapping of the female via the forelegs and proboscis extension [49, 72]. The use of new methods that estimate the full pose of each fly [52] combined with acoustic recordings should address this possibility.

Several recent studies have used hidden state models (HMMs or equivalents) to describe what an animal is doing over time with incredible accuracy [8, 27, 61, 71]. These models take continuous variables (e.g., the angles between an animal’s joints) and discretize them into a set of outputs or behavioral actions. This generates maps of behavioral actions (such as grooming, fast walking, rearing, etc.) and the likelihood of transitions between actions. In this study, the behavior we focus on can also be considered a continuous variable: the male’s song waveform, generated by vibration of his wing. This variable can be discretized into three separate types of song in addition to no song. We show here that it is crucial to sort these actions not simply by the probability of transitioning between them, but according to how feedback cues bias choices between behavioral outputs. In other words, we demonstrate the importance of considering how changes in feedback cues affect the choice of behavioral outputs as well as the transitions between these choices. Animals do not typically switch between behaviors at random, and the GLM of our GLM-HMM provides a solution for determining how feedback cues modulate the choice of behavioral outputs over time. This will be useful not only for the study of natural behaviors, as we illustrate here, but also for identifying when animals switch strategies during task-based behavior [24, 35, 48, 55]. The broader framework presented here can also flexibly incorporate continuous internal states with state-dependent dynamics [46]. Alternately, states themselves may operate along multiple timescales necessitating hierarchical models in which higher-order internal states modulate lower-order internal states, which in turn modulate animal actions [65].

In our study, differences in internal state correspond to differences in how feedback cues pattern song. This is analogous to moving towards someone and engaging in conversation when in a social mood and avoiding eye contact or turning away when not. In both cases, the feedback cues remain the same (the approaching presence of another individual), but what changes is the mapping from sensory input to behavior. Previous studies of internal state have focused mostly on states that can either be controlled by an experimenter (e.g., hunger/satiety) or easily observed (e.g., locomotor status). By using an unsupervised approach to identify states, we expand these studies to states that animals themselves control and are difficult to measure externally. This opens the door to finding the neural basis of these states. We provide an example of this approach by investigating how activation of neurons previously identified to drive song production in *Drosophila* affect the state predictions from the GLM-HMM. We find that activation of a pair of neurons known as pIP10 not only robustly drives the male to produce two types of song (P_fast_ and P_slow_), as shown previously [17], but it also drives males into the ‘close’ state, a state mostly associated with the production of sine, not pulse, song in wild type flies. pIP10 neurons are hypothesized to be postsynaptic to P1 neurons that control the male’s courtship drive [37, 58, 70, 73, 75]. Previous work [19] found that dynamic modulation of pulse song amplitude likely occurred downstream of pIP10, in agreement with what we have found here. In other words, activation of pIP10 neurons both directly drives pulse song production (likely via vPR6, see Fig. 4) and also affects the way feedback cues modulate song choice. While we don’t yet know how this is accomplished, our work suggests pIP10 affects the routing of sensory information into downstream song premotor circuits, analogous to amygdala neurons which both gate sensory information as well as suppress or promote particular behaviors [4, 11, 30, 33].

What insight does our model provide to studies of *Drosophila* courtship more broadly? We expect that internal state also affects the production of other behaviors produced during courtship, such as tapping, licking, orienting, and mounting. This includes not only states like hunger [12, 29, 57, 68], sexual satiety [76], or circadian time [59], but states that change on much faster timescales, as we have observed for acoustic signal generation. Identifying these states will require monitoring feedback cues animals have access to during all behaviors produced during courtship. The feedback cues governing these behaviors may extend beyond the ones described here and may include direct contact between male and female or the dynamics of pheromonal experience. The existence of these states may indicate that traditional ethograms detailing the relative transitions between behaviors exhibit additional complexity, or that there are potentially overlapping ‘state’ ethograms.

Why does the male possess the 3 states that we identified? What is striking about the three states is that feedback cues in one state can have a completely different relationship with song outputs versus in another state. For example, increases in the male’s forward velocity is correlated with increased pulse song in the ‘close’ state but increased sine song in the ‘chasing’ state. These changes in relationship may be due to changing female preferences over time (that is, the female may prefer different types of song at different times, depending on changes in her state), changing goals of the male (potentially, to signal to the female to slow down when she is moving quickly or to prime her for copulation if she is already moving slow), or changes in energetic demands (that is, the male balancing producing the right song for the female with conserving energy). The existence of different states may also generate more variable song over time, which may be more attractive to the female [18, 51], consistent with work in birds [51]. Future studies that investigate the impact of state switching on male courtship success and mating decisions may address some of these hypotheses.

In conclusion, in comparison with classical descriptions of behavior as fixed action patterns [67], even instinctive behaviors like courtship displays are continuously modulated by feedback signals [53]. We show here that, in addition, the relationship between feedback signals and behavior is also not fixed but varies continuously, as animals switch between strategies. Instead, just as feedback signals vary over time, so too do the algorithms that convert these feedback cues into behavior outputs. Our computational models provide a method for estimating these changing strategies, and the essential tools for understanding the origins of variability in behavior.

## Acknowledgements

We thank Minseung Choi, Fred Roemschied, Ramie Fathy, and Dudi Deutsch for assistance with establishing triggered LED stimulation (for optogenetics) in the acoustic behavioral assay chamber. We thank Peter Andolfatto, Vivek Jayaraman, Gerry Rubin, and Barry Dickson for flies. We also thank Georgia Guan for excellent technical assistance, and the entire Murthy lab for feedback on this manuscript. This work was funded by the Simons Foundation SCGB (AJC & JWP; AWD494712, AWD1004351, and AWD543027), NIH BRAIN R01 (MM and JWP), and the Howard Hughes Medical Institute (MM).

## Author Contributions

A.J.C. and M.M. designed the study. A.J.C. collected new data (wild type data analyzed in this study was collected for a previous study [19]). A.J.C. and J.W.P designed the models. A.J.C. analyzed data and generated figures. A.J.C, J.W.P, and M.M. wrote the paper.

## Author Information

Authors have no competing interests. Correspondence and requests for materials should be addressed to.

## Methods

### Flies

For all experiments, we used 3-7 day old virgin flies harvested from density-controlled bottles seeded with 8 males and 8 females. Fly bottles were kept at 25degC and 60% relative humidity. Male virgined flies were then housed individually across all experiments, female virgined flies were group-housed in wild-type experiments and individually housed in the transgenic experiments (Fig 4), and kept in behavioral incubators under 12hr/12hr light/dark cycling. Prior to recording with a female, males were painted with a small spot of opaque ultraviolet-cured glue (Norland Optical and Electronic Adhesives) on the dorsal mesothorax to facilitate identification during tracking. All wild type data in was collected by [19] and consisted of either a random subset of 100 flies not used for model-training (Fig 1g-h, Supplemental Fig. 2c-g) or all wild-type flies in the data set. In Figure 4, we collected additional data using transgenic flies.

**Extended Data Table 1.**
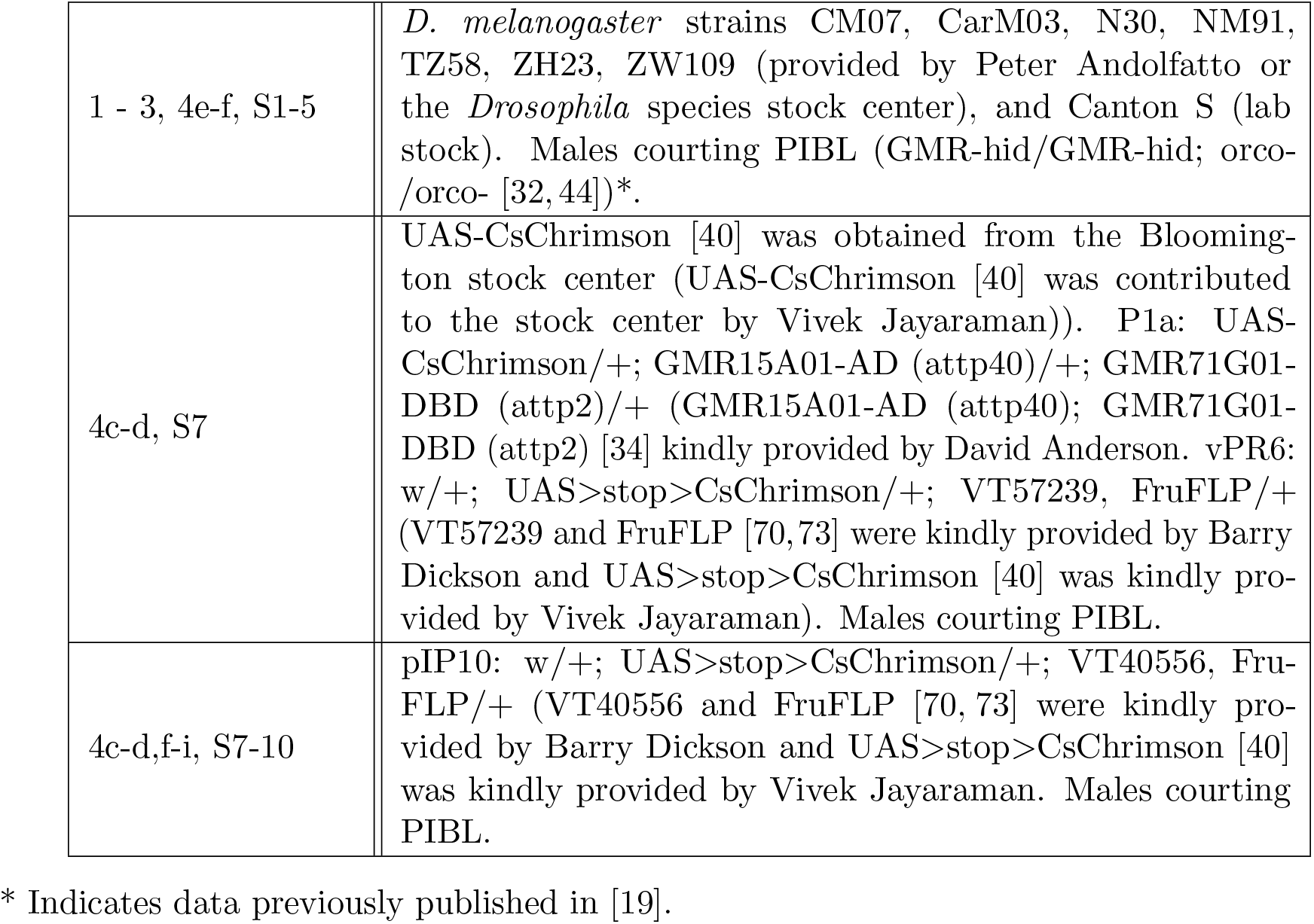

|  |  |
| --- | --- |
| 1 - 3, 4e-f, S1-5 | *D. melanogaster* strains CM07, CarM03, N30, NM91, TZ58, ZH23, ZW109 (provided by Peter Andolfatto or the *Drosophila* species stock center), and Canton S (lab stock). Males courting PIBL (GMR-hid/GMR-hid; orco-/orco- [32,44])*. |
| 4c-d, S7 | UAS-CsChrimson [40] was obtained from the Bloomington stock center (UAS-CsChrimson [40] was contributed to the stock center by Vivek Jayaraman)). P1a: UAS-CsChrimson/+; GMR15A01-AD (attp40)/+; GMR71G01-DBD (attp2)/+ (GMR15A01-AD (attp40); GMR71G01-DBD (attp2) [34] kindly provided by David Anderson. vPR6: w/+; UAS>stop>CsChrimson/+; VT57239, FruFLP/+ (VT57239 and FruFLP [70,73] were kindly provided by Barry Dickson and UAS>stop>CsChrimson [40] was kindly provided by Vivek Jayaraman). Males courting PIBL. |
| 4c-d,f-i, S7-10 | pIP10: w/+; UAS>stop>CsChrimson/+; VT40556, FruFLP/+ (VT40556 and FruFLP [70,73] were kindly provided by Barry Dickson and UAS>stop>CsChrimson [40] was kindly provided by Vivek Jayaraman. Males courting PIBL. |

\* Indicates data previously published in [19].

### Behavioral chamber

Behavioral chambers were constructed as previously described [17, 19, 20]. For optogenetic activation experiments, we used a modified chamber whose floor was lined with white plastic mesh and equipped with 16 recording microphones and video recorded at 60 Hz. To prevent the LED from interfering with the video recording and tracking, we used a short-pass filter (ThorLabs FESH0550, cut-off wavelength: 550 nm). Flies were introduced gently into the chamber using an aspirator. Recordings were timed to be within 150 minutes of the behavioral incubator lights switching on to catch the morning activity peak. Recordings were stopped after 30 minutes or after copulation, whichever was sooner. If males did not sing during the first 5 minutes of the recording, the experiment was discarded.

### Optogenetic activation

Flies were kept for at least 5 days prior to the experiment on either regular fly flood or fly food supplemented with retinal (1ml all-trans retinal solution (100mM in 95% ethanol) per 100ml food). CsChrimson [40] was activated using a 627 nm LED (Luxeon Star) at an intensity of 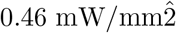 for pIP10 activation. Light stimuli were delivered for 500ms of constant LED illumination, randomized to occur every 5 - 25s. Sound recording and video were synchronized by positioning a red LED that turned on and off with a predetermined temporal pattern in the field-of-view of the camera and whose driving voltage was recorded on the same device as the song recording.

### Fly tracking via DeepFlyTrack

Data from [19] were previously tracked.

For new data, tracking was performed using a custom neural network tracker we call DeepFlyTrack. The tracker has three components: identifying fly centroids, orienting flies, and tracking fly identity across frames. Frames are first annotated to indicate the position of a blinking LED used for synchronization with the acoustic signal and to indicate the portion of the video frame containing the fly arena.

Frames are annotated to indicate the position of a blinking LED used for synchronization with the acoustic signal and to indicate the portion of the video frame containing the fly arena. We designed a neural network trained on 200 frames containing fly bodies annotated with centroid, head, and tail. These annotations were convolved with a 2-dimensional Gaussian with standard deviation of 5. The network was trained to reconstruct this annotated data from grayscale video frames using a categorical cross entropy loss function. The neural network has five 4×4 convolutional layers. The first four layers are passed through a ReLu activation function and the final layer is passed through a sigmoidal nonlinearity. The network was trained using Keras with input frames being a 192×192×3 patch containing 0, 1, or 2 flies. After training, the network predicted whole video frames. These were thresholded and points were fit with k-means, where k=2. To keep track of fly identity, we use the Hungarian algorithm to minimize the distance between flies identified in subsequent frames. The points were fit to an ellipse to extract putative body center and orientation. We use this ellipse for centroid and angle +/− 180. To fully orient flies, we assume that the fly typically moves forward and rarely turns more than 90 degrees per frame. In 1000 frame chunks, we find the 360 degree orientation that best fits these criteria. Position and orientation were smoothed every two frames to downsample from 60 Hz to the 30 Hz used in previous work [19]. Fly identity and orientation were then manually fixed (average 4.5 identity flips/30 minutes).

### Song segmentation

Song data from Coen et al 2014 [19] was re-segmented to separate P_fast_ and P_slow_ according to Clemens et al [17]. New song data (Figure 4) was also segmented using this new pipeline.

### Static Model (‘chance’)

The probability of observing each of the four song modes (“no song”, P_fast_, P_slow_, and “sine”) in a given frame was calculated from a random sample of 40 wild type flies, which we term the ‘static fly’ and denote as *p*_static fly_(song type). We use two ‘Chance’ models: one drawn from song statistics averaged across all of courtship and one drawn from song only at transitions between output types. Thus the probability of observing a particular song mode is determined by:

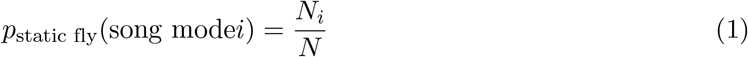

where *N_i_* is the number of time bins during courtship with song mode *i*, and *N* is the total number of time bins, either during all courtship or only at the time of song transitions, averaged across all 40 flies. The likelihood of observing the observed song sequences (Fig 1g-h) was computed using 100 additional flies were sampled from the wild type data set.

### Cross-validation

All hyperparameters were inferred by cross-validation from held-out data not used for assessing performance. Across all analyses, models were fit using one data set and performance was validated on data from individuals that were not used in fitting. Because performance is cross-validated on test data, increasing the number of parameters does not necessarily give higher performance values. See for instance Fig. 1g where the 5-state model achieves lower performance than the 3- or 4-state model while having more free parameters.

### Feedback Cues

Data from tracked fly trajectories were transformed into a set of 17 feedback cues that were considered to affect male singing behavior. For each cue, we extracted 4 seconds of data prior to the current frame, sampled at 30 Hz (120 time samples for each cue), resulting in a feature vector of length 17 × 120 = 2040. We augmented this vector with a ‘1’ to incorporate bias, yielding a vector of length 2041 as input to the model in each time bin.

For model fitting, we formed a design matrix of size *T* × 2041, where *T* is the number of time bins in the dataset from a single fly after discarding the initial four seconds. We concatenated these design matrices across flies so that a single GLM-HMM could be fit to the data from an entire population.

### Multinomial GLM

Previous work [19] used a Bernoulli GLM (also known as logistic regression) to predict song from a subset of the feedback cues that we consider here. That model sought to predict which of two types of song (pulse or sine) a fly would sing at the start of a song bout during certain time windows (e.g., times when the two flies were less than 8 mm apart, and the male had an orientation < 60 degrees from the centroid of the female).

Here we instead use a multinomial GLM (also known as multinomial logistic regression) to predict which of four types of song (no song, P_fast_, P_slow_, and sine) a fly will sing at an arbitrary moment in time. The model is parametrized by a set of four filters {*F*_*S*_}, *S* ∈ {1, 2, 3, 4}, that map the vector of feedback cues to the un-normalized log-probability of each song mode.

The probability of each song mode given under the model given feedback cue vector *s*_*t*_ can be written:

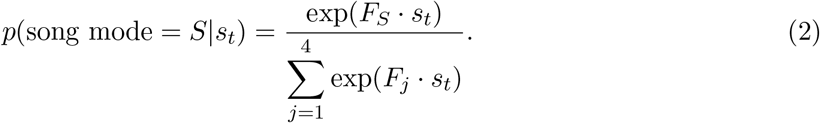

Note that we can set the first filter to all-zeros without loss of generality. We fit the model via gradient descent to maximize the log-likelihood of the observations and used a penalty on the sum of squared differences between adjacent coefficients to impose smoothness. See description of GLM-HMM below for more details (this is a one-state GLM-HMM).

### GLM-HMM

The simplest form of hidden Markov Model (HMM) has discrete hidden states that change according to a set of fixed transition probabilities. At each discrete time step, the model is in one of the hidden states and has a fixed probability of transitioning to another state or staying in the same state. If the outputs are discrete, the HMM has a fixed matrix of emission probabilities, which specifies the probability over the set of possible observations for each hidden state.

The GLM-HMM we introduce in this paper differs from a standard HMM in two ways. First, the probability over observations is parameterized by a GLM, with a distinct GLM for each latent state. This allows for dynamic modulation of output probabilities based on an input vector *s*_*t*_ at each time bin. Second, transition probabilities are also parametrized by GLMs, one for the vector of transitions out of each state. Thus, the probability of transitioning from the current state to another state also depends dynamically on a vector of external inputs (feedback cues) that vary over time.

A similar GLM-HMM was described by Escola et al [26], although it used Poisson GLMs to describe probability distributions over spike train outputs. Here we considered a GLM-HMM with multinomial-GLM outputs, which provides a probability over the four song modes (as described above).

### Fitting

To fit the GLM-HMM to data, we used the expectation-maximization (EM) algorithm [26] to compute maximum-likelihood estimates of the model parameters. EM is an iterative algorithm that converges to a local optimum of the log-likelihood. The log-likelihood (which may also be referred to as the log *marginal* likelihood) is given by:

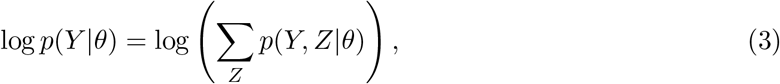

where *Y* = *y*_1_, …, *y*_*T*_ are the observations at each time point and *Z* = *z*_1_, …, *z*_*T*_ are the hidden states that the model enters at each time point. The joint probability distribution over data and latents, known as the “complete-data log-likelihood”, can be written:

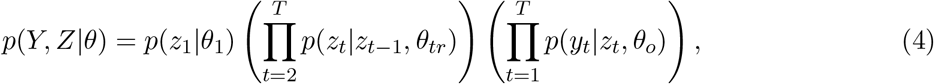

where *θ*_1_ is a parameter vector specifying the probability over the initial latent state *z*_1_, *θ*_*tr*_ denotes the transition model parameters, and *θ*_*o*_ denotes the observation model parameters. We will abbreviate

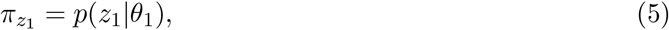

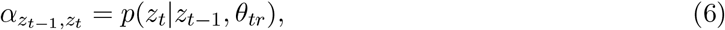

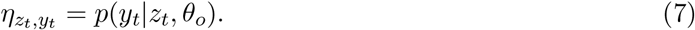

*P*(*Z*_1_|*θ*_1_) is initialized to be uniformly distributed across states and then fit on successive E-steps.

### E-Step

The E-step of the EM algorithm involves computing the posterior distribution *p*(*Z*|*Y,θ*) over the hidden variables given the data and model parameters. We use the adapted version of the Baum-Welch algorithm described in Escola et al [26]. The Baum-Welch algorithm has two components, a forward step and a backward step. The forward step identifies the probability *a*_*m*_(*t*) = *p*(*Y*_1_ = *y*_1_, …, *Y*_*t*_ = *y*_*t*_, *Z*_*t*_ = *m*|*θ*) of observing *Y* = *y*_1_, *y*_2_, …, *y*_*t*_ and, assuming there are *N* total states, of being in state *m* at time *t* by iteratively computing:

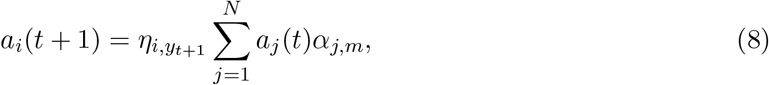

where *a*_*i*_(1) = *π*_*i*_. The backward step does the reverse—it identifies the conditional probability of future observations given the latent state:

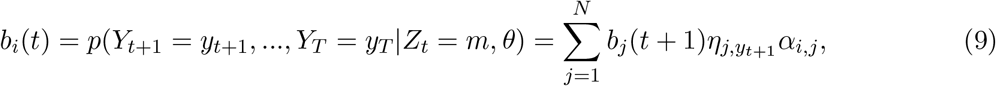

where *b*_*i*_(*T*) = 1. These allow us to compute the marginal posterior distribution over latent state at every time step, which we denote *γ*(*Z*_*t*_) = *P*(*Z*_*t*_|*Y,θ*), and over pairs of adjacent latent states, denoted *ξ*(*Z*_*t*_, *Z*_*t*+1_) = *P*(*Z*_*t*_, *Z*_*t*+1_|*Y,θ*), which are given by:

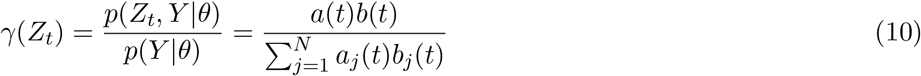

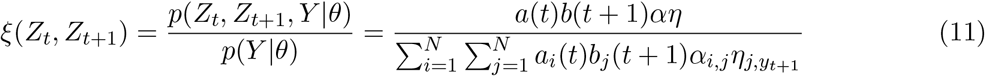

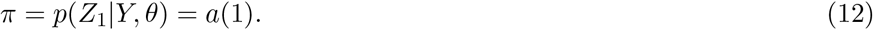

In practice, for larger data sets it is common to run into underflow errors due to repeated multiplication of small probabilities in the equations above. Thus, it is typical to compute {*a*_*m*_(*t*)} and {*b*_*m*_(*t*)} in scaled form. See [54] for details.

### M-Step

The M-step of the EM algorithm involves maximizing the expected complete-data log-likelihood for the model parameters, where expectation is with respect to the distribution over latents computed during the E-step:

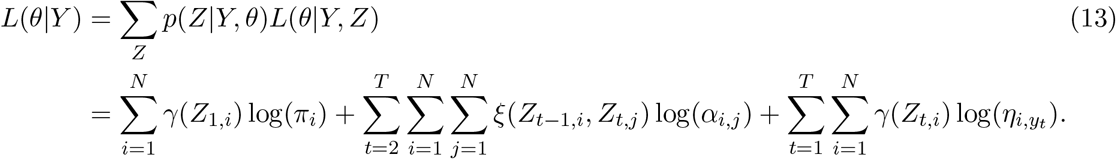

As noted above, our model describes transition and emission probabilities with multinomial GLMs, each of which is parametrized by a set of filters. Because these GLMs contribute independently to *α*, *η*, and *π* terms above, we can optimize the filters for each model separately. Maximizing the *π* term is equivalent to finding *π*_*i*_ = *p*(*Z*_1_ = *i*|*Y,θ*). We maximize the *α* term as described in Escola et al (Appendix B) with the addition of a regularization parameter. The transition probability from state *i* to state *j* at time *t* is defined to be

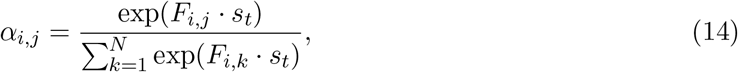

where we define all filters from one state to itself *F*_*i,i*_ to be 0 without loss of generality.

Additionally, we have added regularization penalties into the model in order to avoid overfitting. We tried using both Tikhonov regularization and difference smoothing and found difference smoothing to provide both better out-of-sample performance and filters that were less noisy. Difference smoothing adds a penalty for large differences in adjacent bins in each filter. However, because each filter was applied across *U* features of length *L* we did not apply a penalty between bins across features. For some regularization coefficient *r*, the model that we fit became:

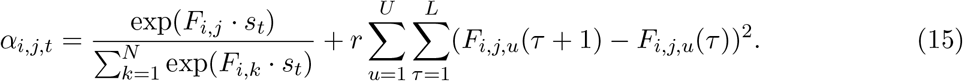

Escola et al [26] provides both the gradient and Hessian for fitting the transition filters during the M-step, although in our hands we found computing the Hessian to be computationally more expensive for the large data sets that we are working with and does not speed up the fitting procedure. We computed the inverse Hessian at the end of each stage of fitting to provide an estimate of the standard error of the fit.

The GLM-HMM described by Escola et al [26] was formulated for neural data, in which the outputs at each time were Poisson or Bernoulli random variables (binned spike counts). As noted above, we have modified the model to use categorical outputs to predict the discrete behaviors the animal is performing. Similar to the transition filters, for emission filters one filter may be chosen to be the ‘baseline’ filter set to 0. We used a multinomial model which assumes coefficients *F* = *F*_1_, …, *F*_*n*_ where the filter *F*_1_ is assumed to be the ‘baseline’ filter equal to 0. The probability of observing some output *y*_*t*_ (out of *O* possible outputs) is governed by the equation

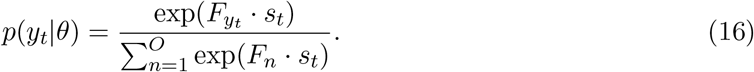

We then maximize the following:

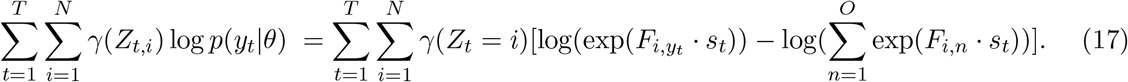

In order to avoid overfitting, we increased smoothness between bins by penalizing differences between subsequent bins for each feature using the difference operator *D* multiplied by the regularization coefficient *r* as described for the transition filter. No smoothing penalty was applied at the boundary between features. The objective function to optimize becomes:

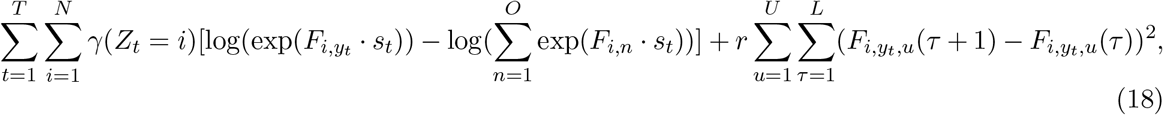

which has gradient

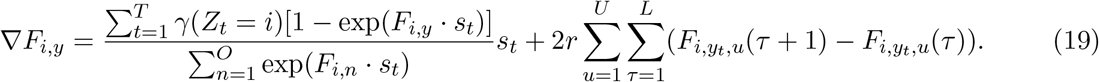

After the M-step is complete, we ran the E-step to find the new posterior given the parameters found in the M-step, and continued alternating E and M steps until the log-likelihood increases by less than some threshold amount.

We used the minfunc function in MATLAB to minimize the negative expected complete-data log-likelihood in the M step, and used cross-validation to select regularization penalty *r* over a grid of values. The selected penalty for the 3-state model was *r* = 0.05.

### Testing

We assessed model performance by calculating the log-likelihood of data held out from training. In particular, we assessed how well the model can predict the next output given knowledge of all the data up to the present moment. This allows for an accurate estimate of the state up to time *t* and then evaluates how well the transition and output filters explain what happens at time *t* + 1. We can write this as log(*p*(*Y*_*t*+1_|*Y*_1_ = *y*_1_, …, *Y*_*t*_,*θ*)) and can be calculated in the same manner as the forward pass of the Expectation step described in the previous section. The prediction for *t* + 1 is then

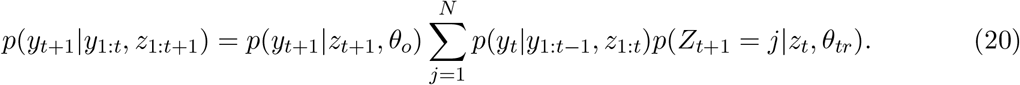

Note that this equivalent to the scaling factor used when fitting the forward step of the model in the preceding section. The mean forward log-likelihood that we report is then

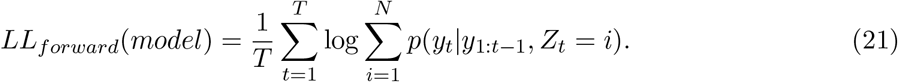

We normalized the forward log-likelihood by the performance of the ‘Chance’ model (described above, Fig. 1g, Supplemental Fig. 2c,e),

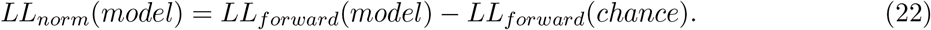

The ‘Chance’ model was either drawn from the entirety of courtship (Fig. 1g) or only at transitions between outputs (Fig. 1h) (see Static model ‘chance’, above). While this provides a bound on the level of uncertainty of each model, we do not have a way of estimating the intrinsic variability in the male’s song patterning system (to do so, would require presenting the identical feedback cue history twice in a row, with the animal in the same set of states).

An alternative metric that we report is the log-likelihood of the model where we did not use the past song output to estimate the current state. This alternative requires us to only use the state transition filters to predict current state. Recall from the previous section that *p*(*z*_*t*_|*z*_*t*−1_, *θ*_*tr*_) is the probability of transitioning using only the transition model. Then the probability of being in each state at time *t* is:

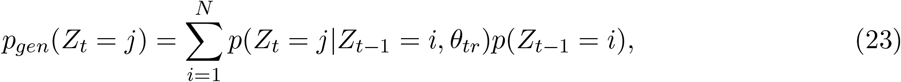

and the log-likelihood of the output is

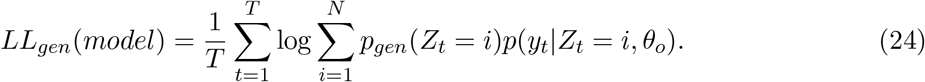

Giving a normalized log-likelihood measure similar to before:

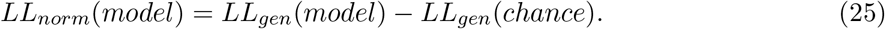

The normalized log-likelihood of the forward model reports improvement over the ‘Chance’ model for predicting song mode given knowledge of song history to improve the prediction of the current state. The normalized log-likelihood using only the feedback cues reports improvement over the ‘Chance’ model for predicting song mode, without accurately estimating the state of the animal in other words, model performance that comes only from dynamics of the feedback cues.

### Binning of song

We discretized the acoustic recording data to fit the GLM-HMM. Song was recorded at a sampling rate of 10KHz, segmented into 4 song modes (sine, P_fast_, P_slow_, and no song), and then discretized into time bins of uniform width (33 ms) which corresponds to roughly the inter-event interval between pulse events. We use the modal type of song in each bin to define the song mode in that bin. Because some song pulses have an inter-event interval of > 33ms, we artificially introduced “no song” bins within trains of pulses. To correct this error, we identified the start and end of a run of song as either a transition between types of song (sine, P_fast_, or P_slow_) or as the transition between a song type and “no song”, if the quiet period lasted for > than 80ms. We then corrected “no song” bins that occurred within runs of each song type. In order to up- and down-sample song (Supplemental Fig. 2a,c), we use the modal song per interpolated bin.

### Applying filters from only one state (Fig 3A-B)

The likelihood of observing song is typically calculated by applying the filters for each state and multiplying that by the probability of being in each of those states *p*(state|data)*p*(emission, state). To calculate the probability of being in a given state, we use the Viterbi algorithm used in HMMs to find the most likely state. We then apply the filter for state *i* when the most likely state at time *t* is *j* to find the mean likelihood of observing an emission *k*.

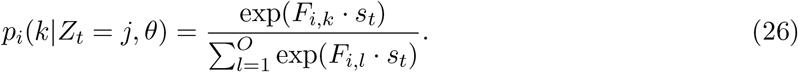

### Predictions of pulse versus sine (Fig 3E)

When fitting multinomial GLM filters, one set of filters is set to 0 and is used as a reference point for other filters (since it is always possible to scale all filters together). In our model, the ‘no song’ output is the filter set to 0. To visualize the filter that represents the probability of observing pulse versus sine, we take the average of the P_fast_ and P_slow_ filters and subtract that from the sine filter.

We then compute the raw data by taking the histogram of Z-scored feedback cues that occur just prior to pulse and divide it by the histogram of Z-scored feedback cues that occur just prior to both pulse and sine. This gives us the probability of observing pulse versus sine at each feature value.

### Fit of GLM-HMM to optogenetic activation (Fig 4)

Optogenetic activation (driven by an LED stimulus) of previously identified song pathway neurons can, as expected, directly drive song production. To account for this, we add in an additional offset term (because each genotype produces different distributions of types of song) and an extra filter for the LED (that is, the LED stimulus pattern is another input to the model, similar to the 17 feedback cues), and then refit the GLM-HMM. The offset term and the additional LED filter are fit using expectation maximization (see above); no other filters (for the 17 feedback cues) are refit from the original GLM-HMM.

### Normalization of pulse/sine ratio (Fig 4)

Optogenetic activation of courtship neurons dramatically changes the fixed probability of observing pulse song. In order to visualize the song output during this activation in spite of the decreased dynamic range, we normalize the output by subtracting the minimal probability of observing pulse vs sine song and dividing by the maximum. This keeps the shape and relationship of the pulse vs sine data constant but compresses it to be between 0 and 1.

### Comparison of GLM-HMM to Coen et al 2014 (Supplemental Fig. 1b)

Coen et al. pCorr values were taken from that paper. In order to generate a fair comparison, we took equivalent song events in our data (for example, ‘pulse start’ compared times at which the male was close to and oriented towards the female and either started a song bout in pulse mode, or did not start a song bout) and found (with either the multinomial GLM or the 3-state GLM-HMM) the song mode (‘pulse’, ‘sine’, or ‘no song’) that generated the maximum likelihood value. Similar to Coen et al., we adjusted the sampling of song events to calculate pCorr. P_fast_ and P_slow_ events were both counted as ‘pulse’. The pCorr values reported for the multinomial GLM and 3-state GLM-HMM are the percent of time the highest likelihood value corresponded to the actual song event.

### Zeroing out filters for Fig 4

To assess performance of the model without key features (male feedback cues or female feedback cues), the filters were set to 0. This removes the ability of the model to perform any prediction with this feature. We then inferred the likelihood of the data using this new model.

## Data availability

Data available at http://arks.princeton.edu/ark:/88435/dsp01rv042w888.

## Code availability

Code for tracking flies (“DeepFlyTrack”) is available at https://github.com/murthylab/DeepFlyTrack. Code for running the GLM-HMM algorithm is available at https://github.com/murthylab/GLMHMM.

## Supplementary Information

**Figure S1.**
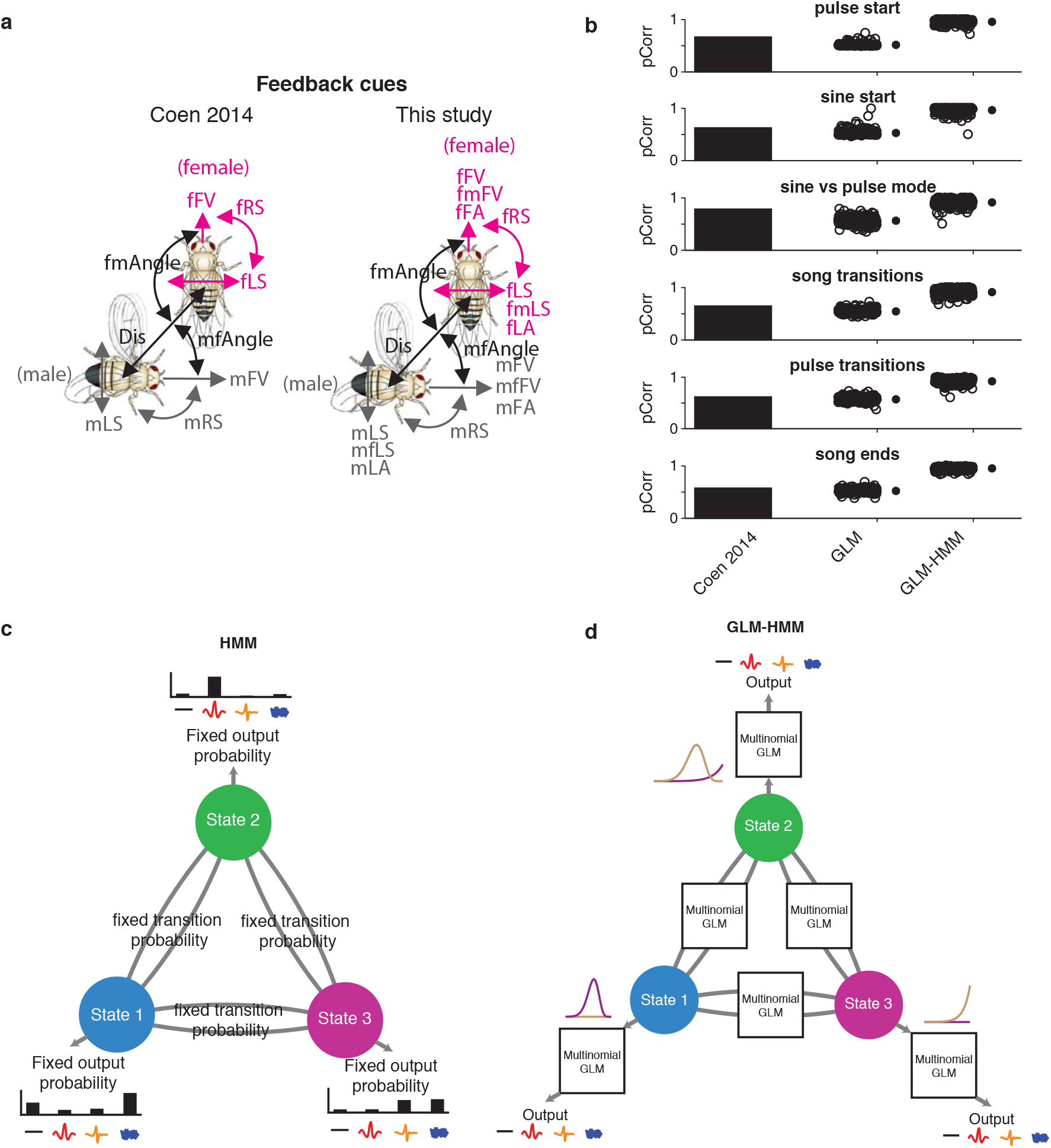
**a.** Fly feedback cues used for prediction in (Coen et al. 2014) (left) or the current study (right). **b.** Comparison of model performance using probability correct (‘pCorr’) (see Methods) for predictions from (Coen et al. 2014) (reproduced from that paper) for the single-state GLM (See Fig. 1c) and 3-state GLM-HMM (see Fig. 1d). Each open circle represents predictions from one courtship pair. The same pairs were used when calculating the pCorr value for each condition (GLM and 3-state GLM-HMM); filled circles represent mean +/− SD; 100 shown for visualization purposes. **c.** Schematic of standard HMM, which has fixed transition and emission probabilities. **d.** Schematic of GLM-HMM in the same format, with static probabilities replaced by dynamic ones. Example filters from the GLM are indicated with the purple and brown lines.

**Figure S2.**
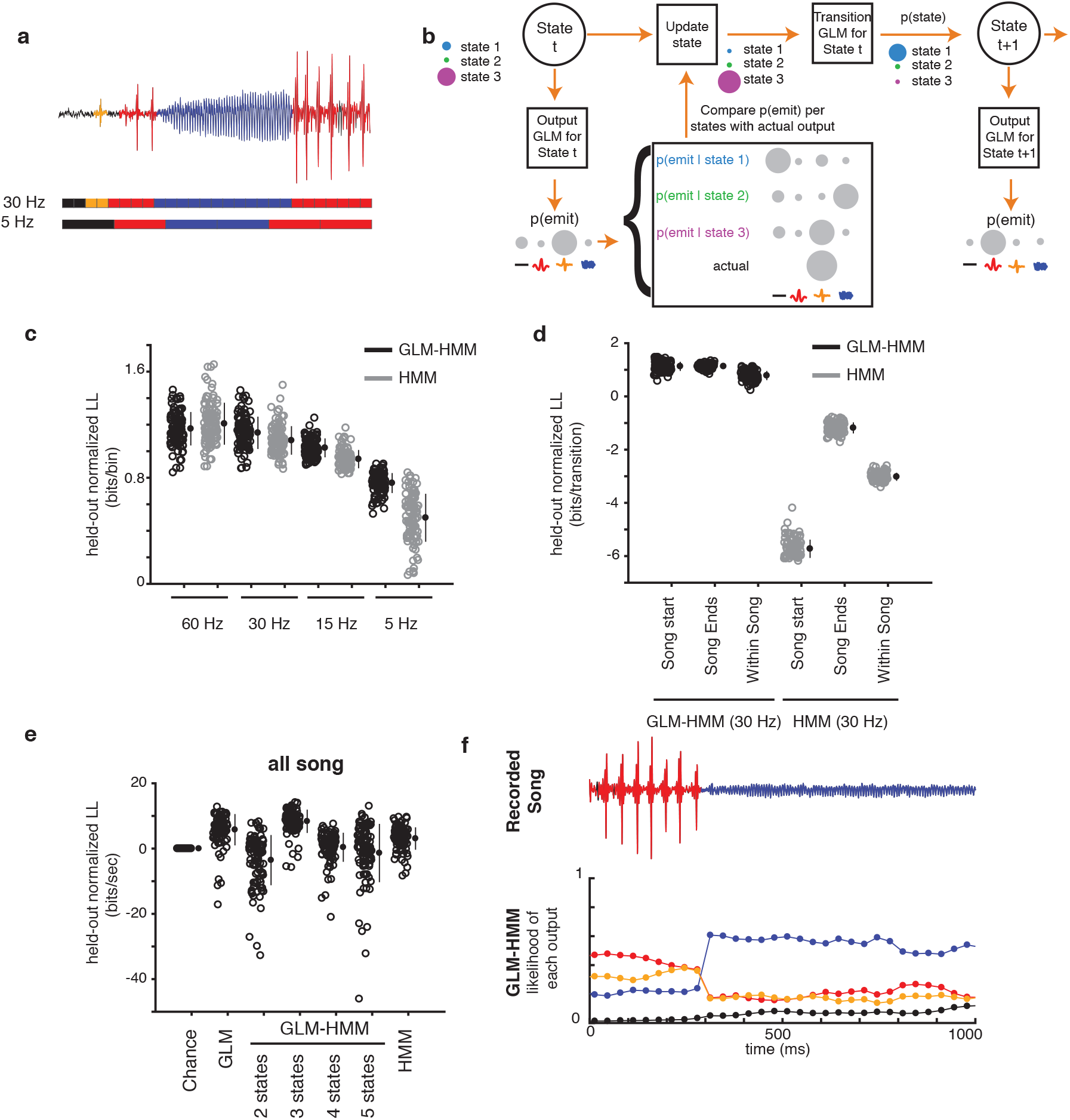
**a.** Illustration of how song is binned for model predictions. Song traces (top) are discretized by identifying the most common type of song in between two moments in time, allowing for either fine (middle) or coarse (bottom) binning - see Methods. **b.** Illustration of how model performance is estimated, using one step forward predictions (see Methods). **c.** 3-state GLM-HMM performance (in bits/sec) when song is discretized or binned at different frequencies (60 Hz, 30 Hz, 15 Hz, 5 Hz) and compared to a static HMM - all values normalized to a ‘Chance’ model (see Methods). Each open circle represents predictions from one courtship pair. Filled circles represent mean +/− SD, n=100. **d.** Comparison of the 3-state GLM-HMM with a static HMM for specific types of transitions when song is sampled at 30 Hz (in bits/transition) - all values normalized to a ‘Chance’ model (see Methods). The HMM is worse than the ‘Chance’ model at predicting transitions. Filled circles represent mean +/− SD, n=100. **e.** Performance of models when the underlying states used for prediction are estimated ignoring past song mode history (see b) and only using the the GLM filters - all values normalized to a ‘Chance’ model (see Methods). The 3-state GLM-HMM significantly improves prediction over ‘Chance’ (p ¡ 0.01, t test) and outperforms all other models. Filled circles represent mean +/− SD, n=100. **f.** Example output of GLM-HMM model when the underlying states are generated purely from feedback cues (e).

**Figure S3.**
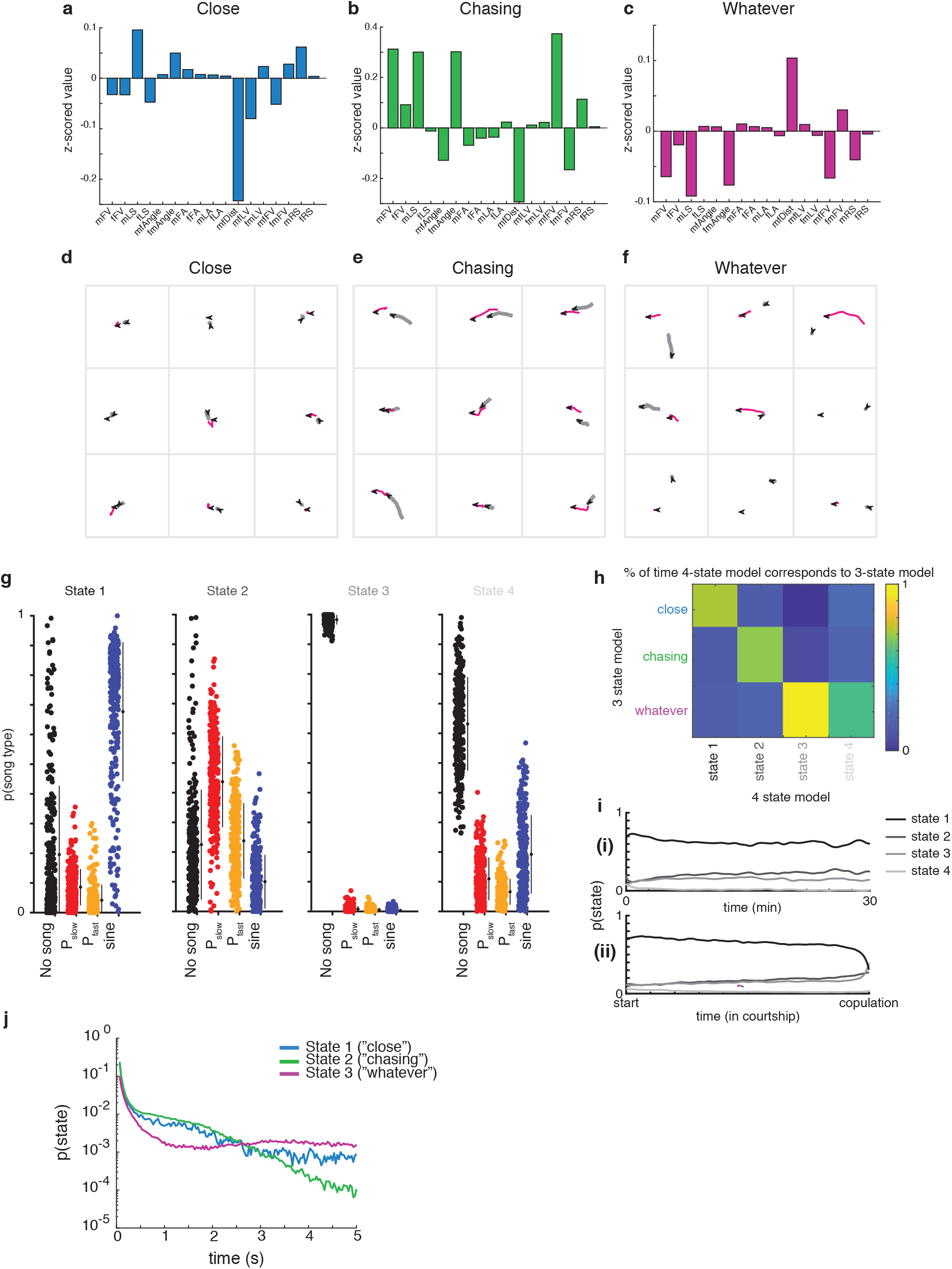
**a-c.** The mean value for each feedback cue in the **(a)** ‘close’, **(b)** ‘chasing’, or **(c)** ‘whatever’ state (see Methods for details on z-scoring). **d-f.** Representative traces of male and female movement trajectories in each state. Male trajectories are in gray and female trajectories in magenta. Arrows indicate fly orientation at the end of 660 ms. **g.** In the 4-state GLM-HMM model, the probability of observing each type of song when the animal is in that state. Filled circles represent individual animals (n=276 animals, small black circles with lines are mean +/− SD). **h.** The correspondence between the 3-state GLM-HMM and the 4-state GLM-HMM. Shown is the conditional probability of the 3-state model being in the ‘close’, ‘chasing’, or ‘whatever’ states given the state of the 4-state model. **i.** The mean probability across flies of being in each state of the 4-state model when aligned to absolute time (top) or the time of copulation (bottom). **j.** Probability of state dwell times generated from feedback cues. These show non-exponential dwell times on a y-log plot.

**Figure S4.**
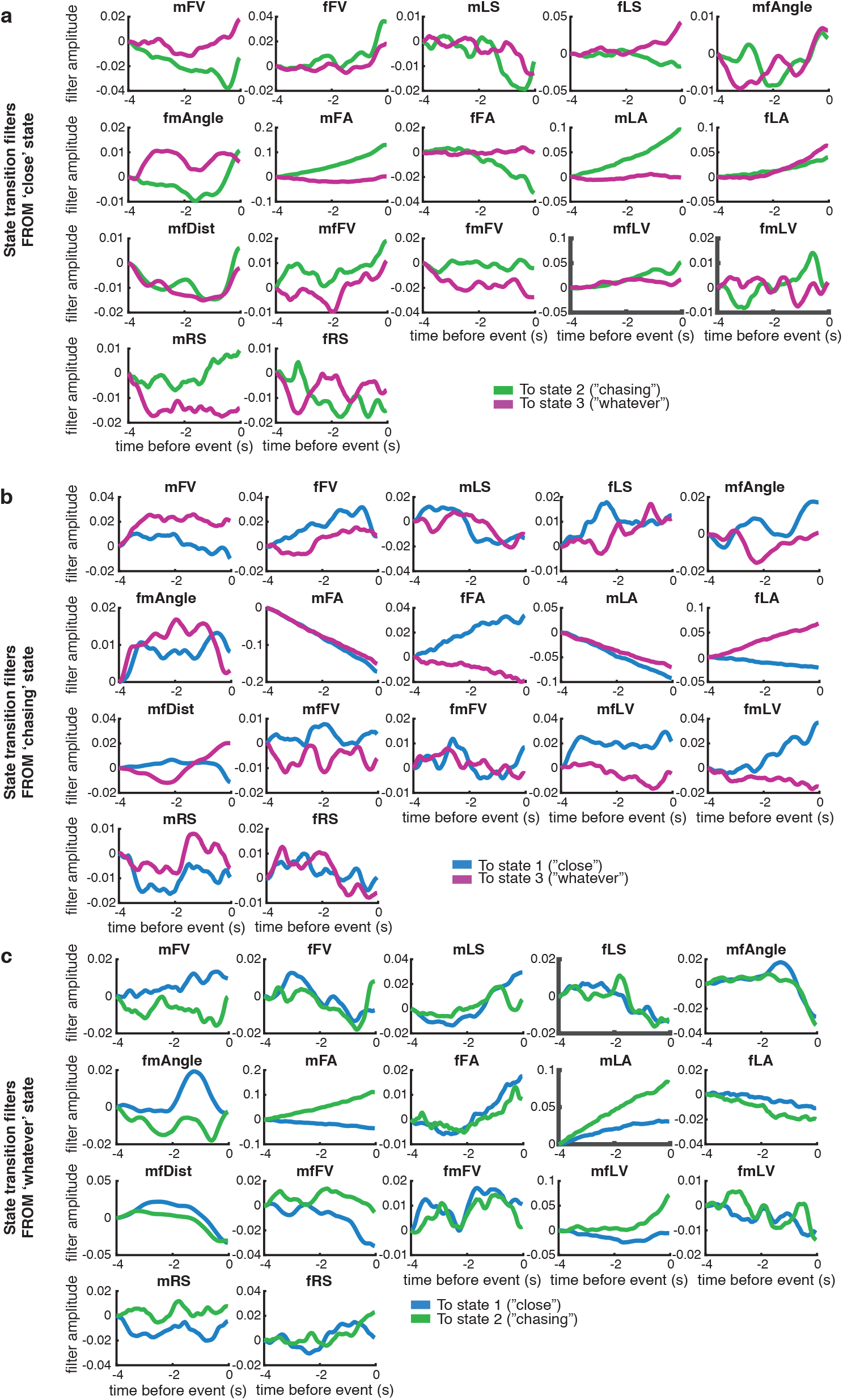
**a-c.** State-transition filters that predict transitions from one state to another for each feedback cue (see Fig. 1B for list of all 17 feedback cues used in this study).

**Figure S5.**
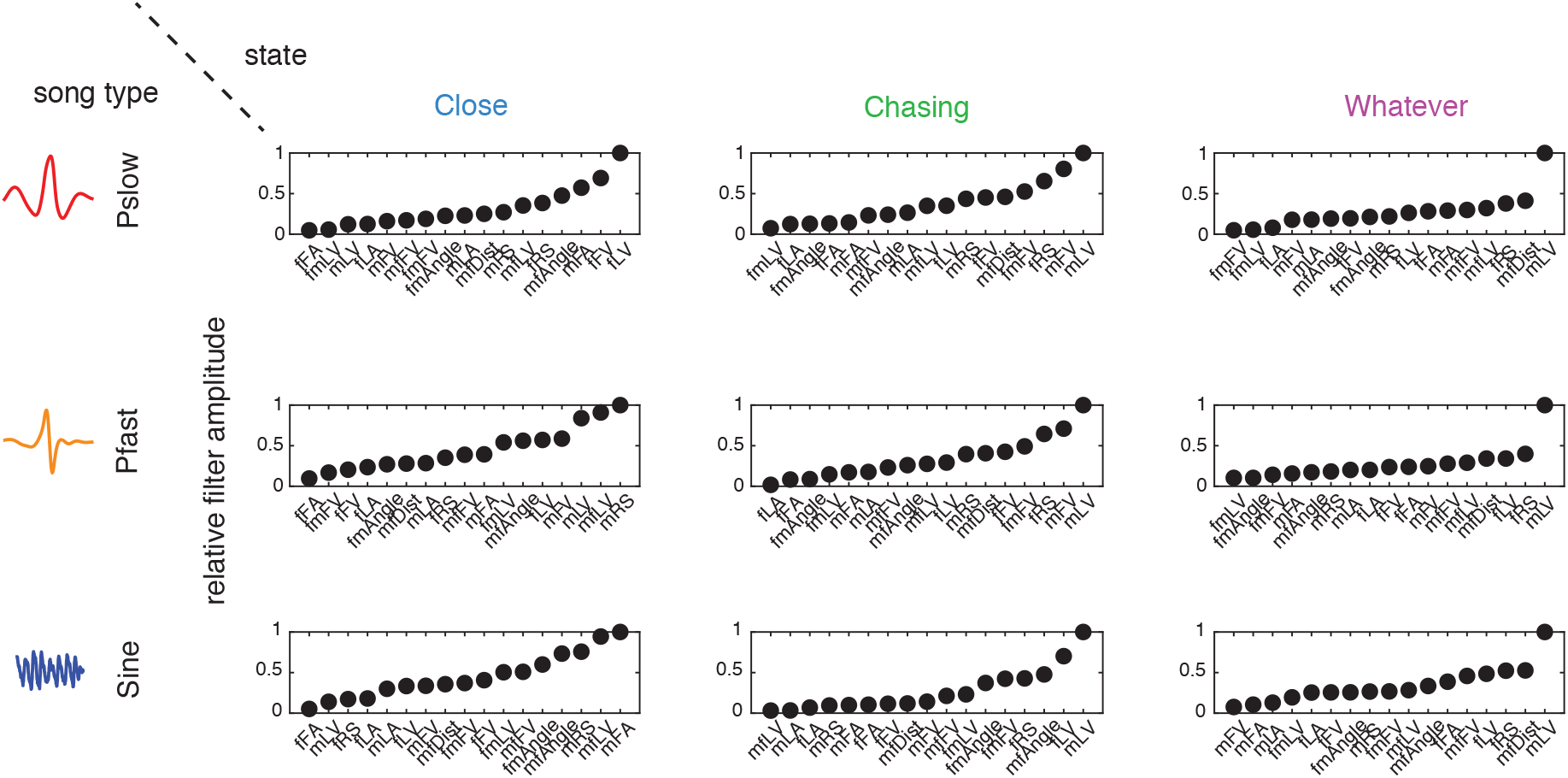
The amplitude of output filters (see Methods) for each state/output pair. Output filter amplitudes were normalized between 0 (smallest filter amplitude) and 1 (largest filter amplitude).

**Figure S6.**
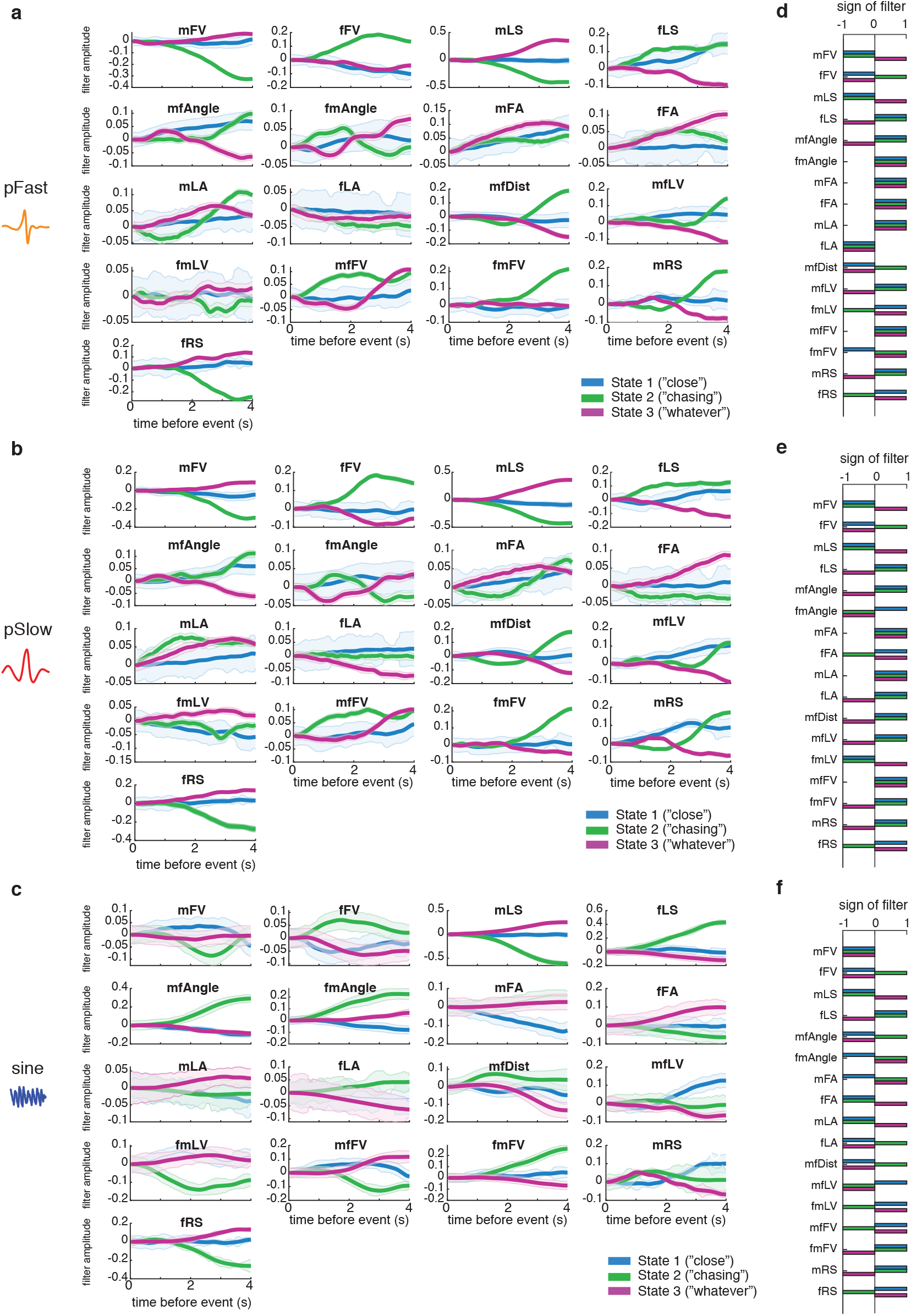
**a-c.** Output filters for each feedback cue (see Fig. 1b) that predict the emission of each song type for a given state. ‘No song’ filters are not shown as these are fixed to be constant, and song type filters are in relation to these values (see Methods). **d-e.** Sign of filter for each emission filter shows the same feature can be excitatory or inhibitory depending on the state.

**Figure S7.**
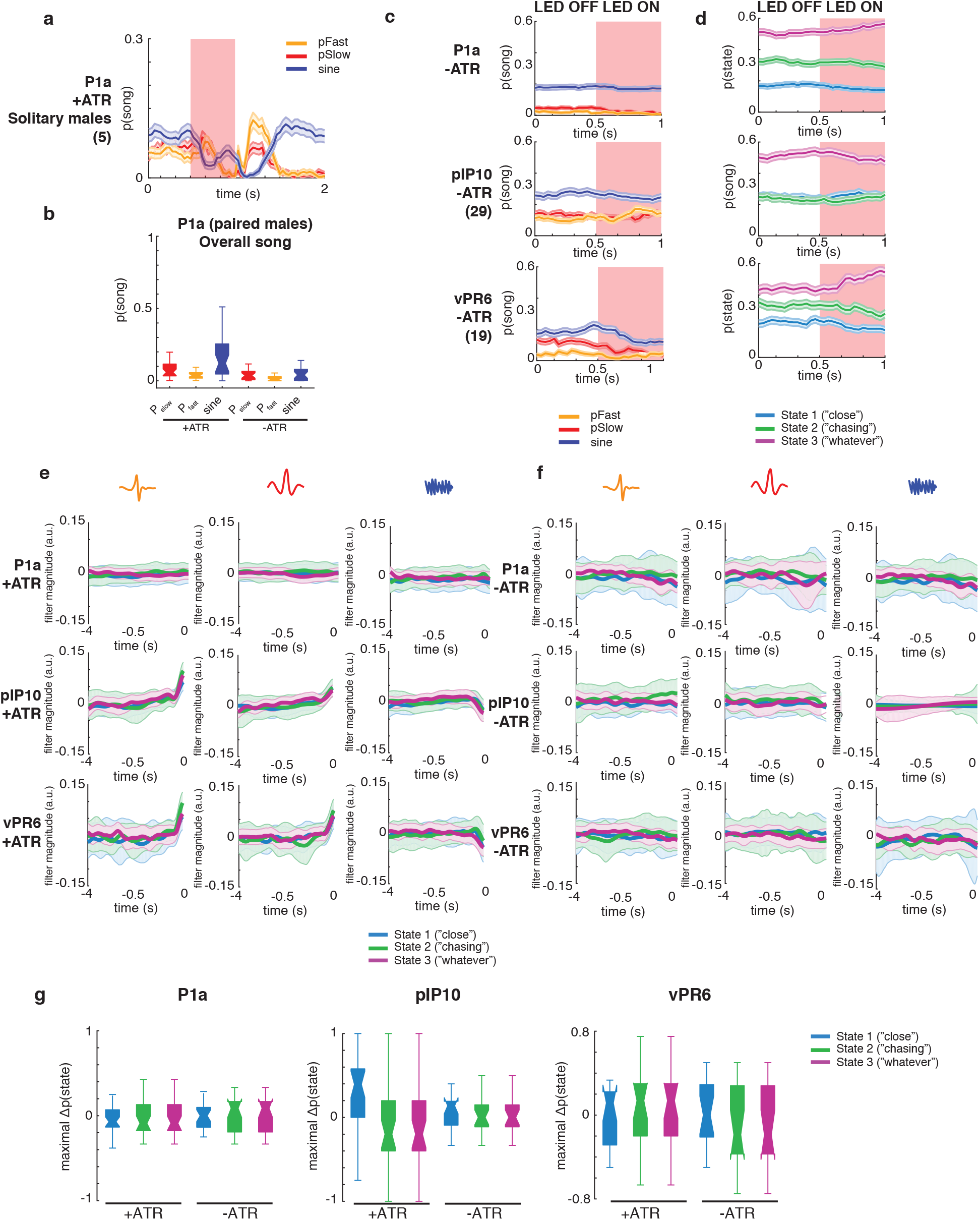
**a.** Solitary ATR-fed P1a males produce song when exposed to the same LED stimulus used in Figure 4. In solitary males, song production is both long-lasting and time-locked to the LED stimulus. **b.** ATR-fed P1a males courting a female produce significantly more (P_fast_) and (P_slow_) (p ¡ 0.01) across courtship but not significantly-different amounts of sine song (p = 0.5). Center lines of box plots represent median, the bottom and top edges represent the 25th and 75th percentiles (respectively). Whiskers extend to +/− 2.7 times the standard deviation. All p-values from two-tailed t test. **c.** The probability of observing each song mode aligned to the opto stimulus shows that LED activation of flies not fed ATR does not increase song production. **d.** The probability of the model being in each state aligned to the opto stimulus shows that LED activation of flies not fed ATR does not change state residence. Error bars represent SEM. **e-f.** ‘Opto’ filters represent the contribution of the LED to the production of each type of song for (e) ATR+ and (f) ATR− flies. The filters for each strain and song type are not significantly different between states. **g.** Measuring the maximal change in state probability between LED ON and LED OFF shows that only pIP10 activation produces a significant difference between ATR+ and ATR− flies (two-tailed t test).

**Figure S8.**
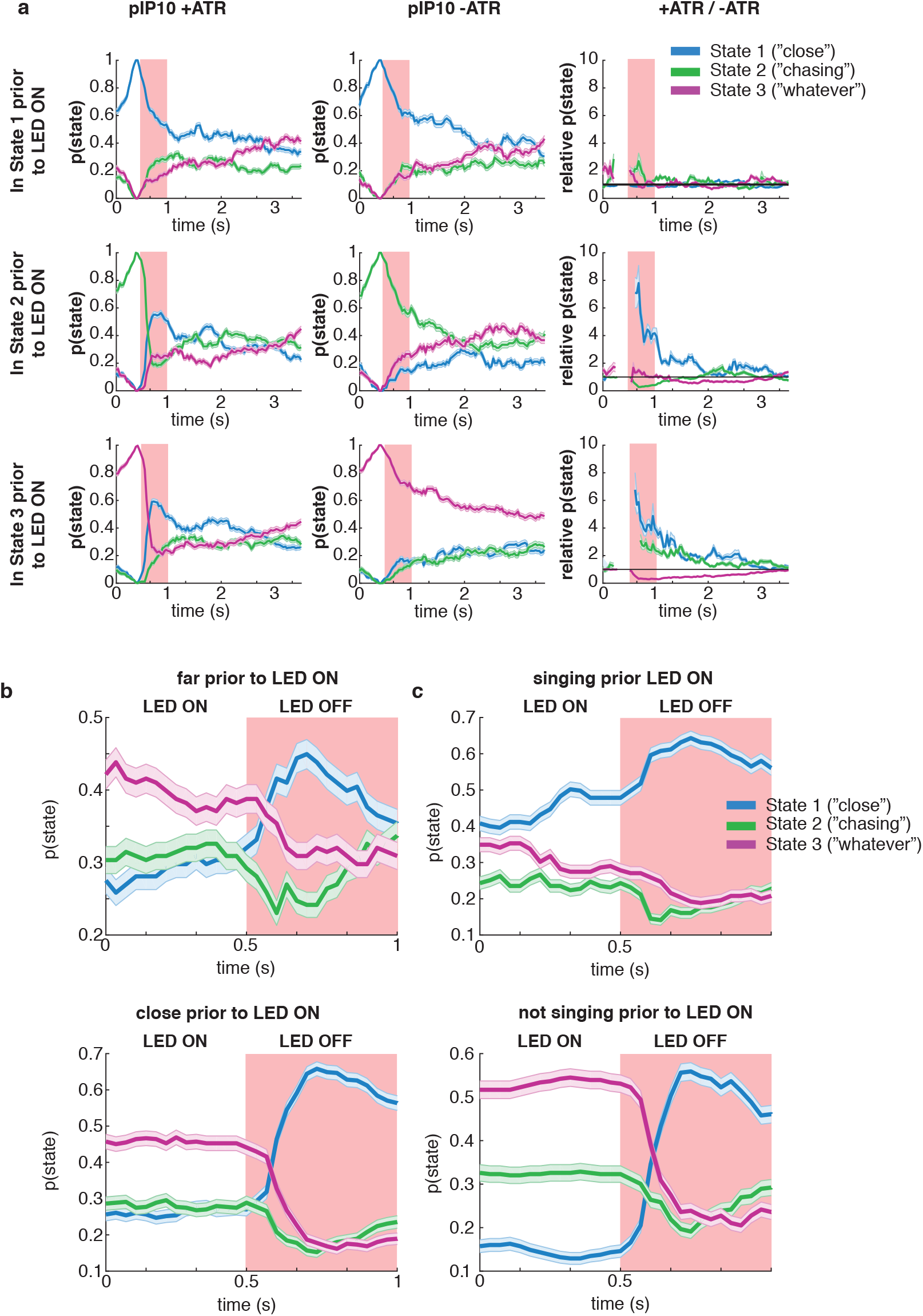
**a.** Conditioning on which state the animal is in prior to the light being on (left, ATR-fed pIP10 flies; middle ATR-free pIP10 flies; right, ratio of ATR-fed to ATR-free state dwell time), activation of pIP10 results in an increase in the probability of being in the close state unless the animal was already in the close state. Shaded area is SEM. **b.** When the male was both close (<5mm) and far (> 8mm), pIP10 activation increases the probability that the animal will enter the close state. Shaded area is SEM. **c.** When the male was already either singing or not singing, pIP10 activation increases the probability that the animal will enter the close state. Shaded area is SEM.

**Figure S9.**
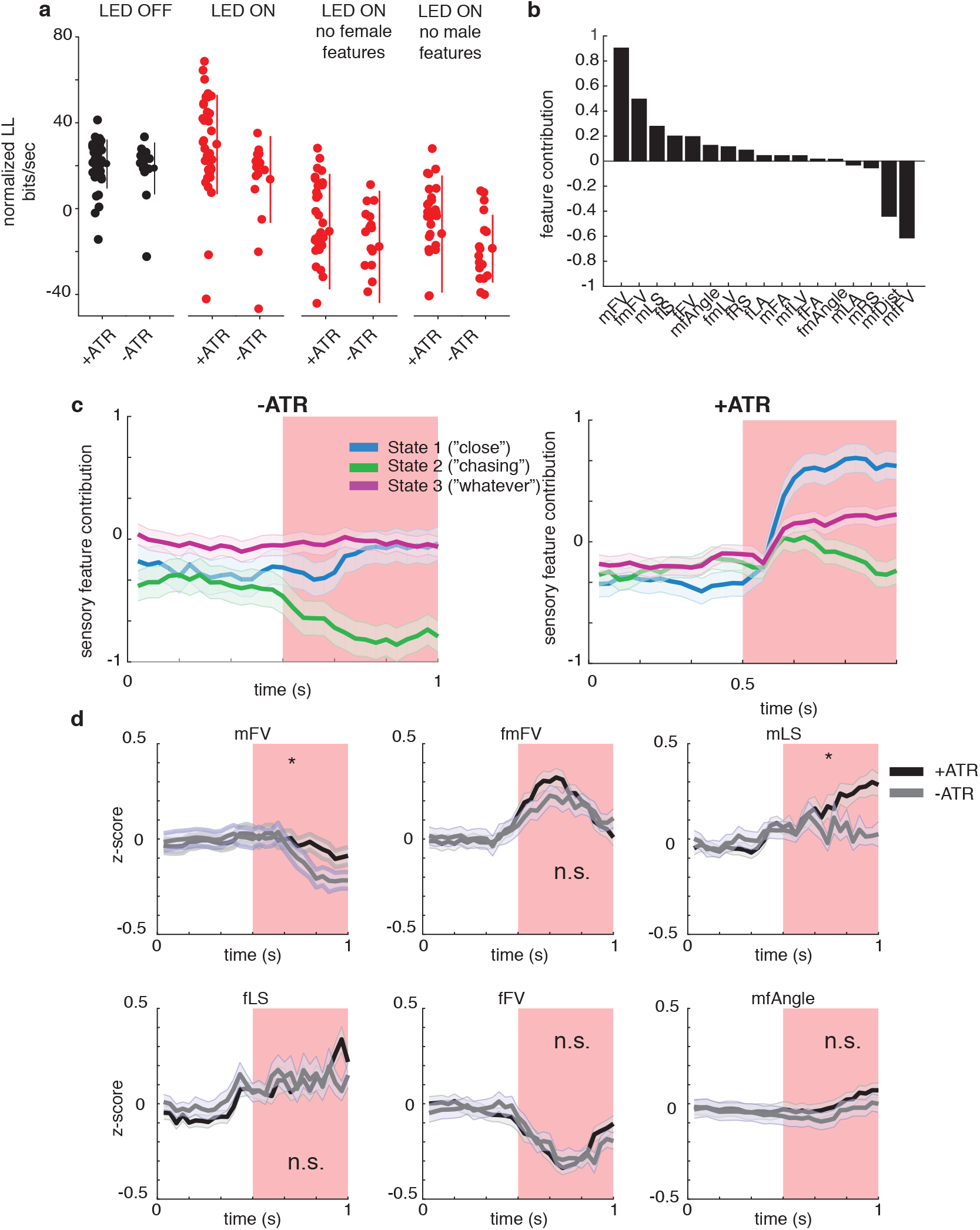
**a.** Predictive performance is not significantly different between light ON and light OFF conditions for both ATR-fed and ATR-free animals (p > 0.2, two-sample t test). Performance suffers without male or female feedback cues, suggesting these state-specific features are needed to predict animal behavior (p < 1e-10, two-sample t test corrected for multiple comparisons with Bonferroni correction). Dots represent individual flies and lines are +/− SD. **b.** The similarity between each feedback cue and the filters for the ‘close’ state are subtracted by the similarity of that feedback cue to the filters for the ‘chasing’ state during LED activation of ATR-fed pIP10 flies. This reveals song patterning is more similar to the ‘close’ state than the ‘chasing’ state for most feedback cues. **c.** Animals that were not fed ATR do not show a change in the contribution of the feedback cues to being in a given state, while animals that are fed ATR do show a change in feedback cue contribution. Shaded area is SEM. **d.** Most aspects of the animal trajectory do not differ in response to red light when males are either fed (black) or not fed (gray) ATR food. Plotted are the six strongest contributors from (b). p-values from two-tailed t test; * represents p < 0.05, n.s. p > 0.05. Shaded area is SEM.

**Figure S10.**
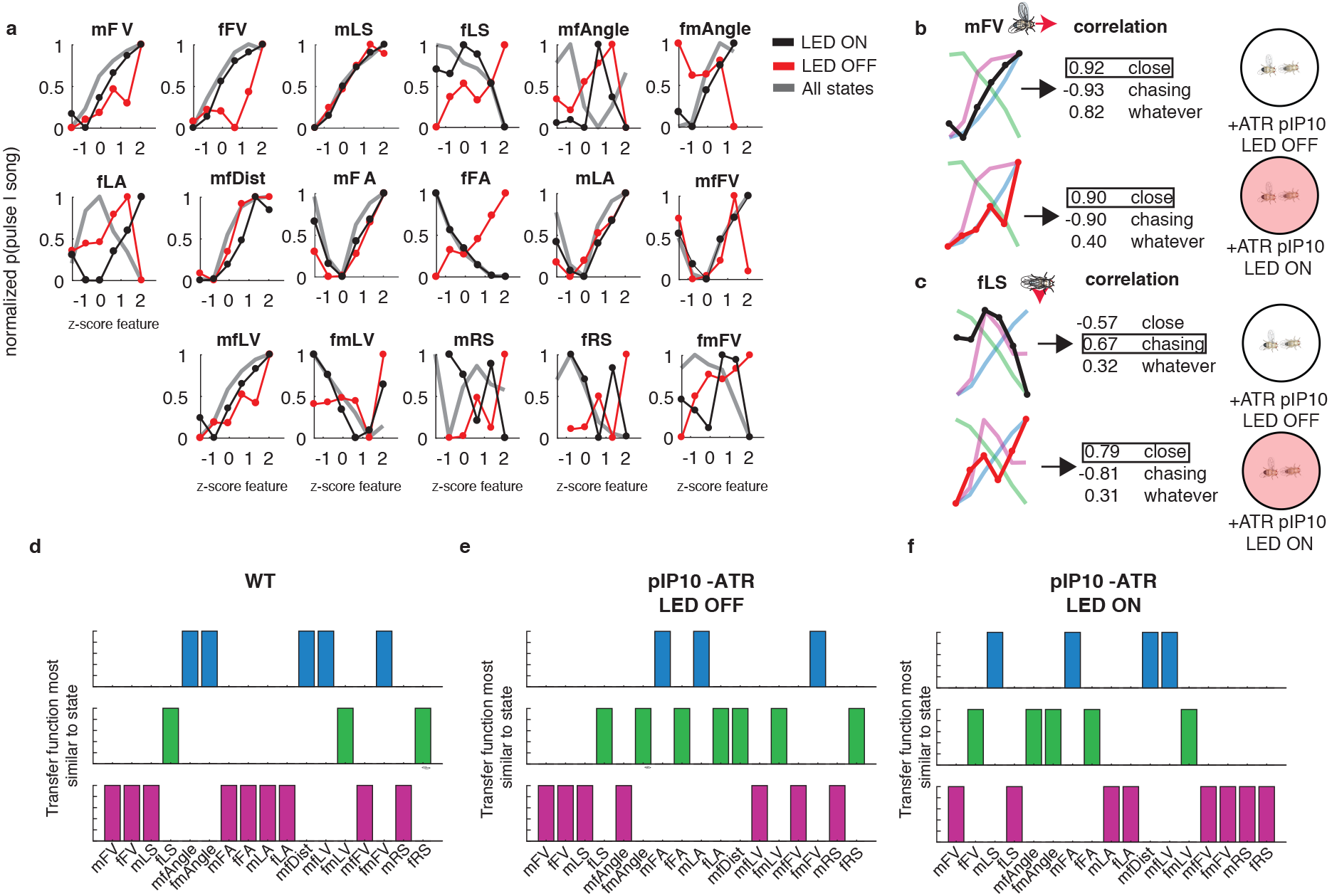
**a.** The transfer functions of each feedback cue when the LED is OFF (black) and when the LED is ON (red) compared to the wild-type average (dark gray). **b-c.** Illustration of correlation between transfer functions of two feedback cues (mFV (g) and fLS (h)) when the LED is off (top, black) and on (bottom, red) and the transfer function in each state (not the average as in Fig. 4f). The feature is considered most similar to the state with which it has the highest correlation. **d.** The state transfer function that is closest to the wild-type average for each feedback cue. For instance, the mFV average is closest to the ‘whatever’ state and the fLS average is closest to the ‘chasing’ state. **e-f.** Same as (d), but for pIP10 flies that not fed ATR food when the LED is off (e) or on (f).

